# Cell cycle exit and stem cell differentiation are coupled through regulation of mitochondrial activity in the Drosophila testis

**DOI:** 10.1101/2021.01.12.426342

**Authors:** Diego Sainz de la Maza, Silvana Hof-Michel, Lee Phillimore, Christian Bökel, Marc Amoyel

## Abstract

Stem cells maintain tissue homeostasis by proliferating to replace cells lost to damage or natural turnover. Whereas stem and progenitor cells proliferate, fully differentiated cells exit the cell cycle. How cell identity and cell cycle state are coordinated during this process is still poorly understood. The Drosophila testis niche supports germline stem cells and somatic cyst stem cells (CySCs), which are the only proliferating somatic cells in the testis. CySCs give rise to post-mitotic cyst cells and therefore provide a tractable model to ask how stem cell identity is linked to proliferation. We show that the G1/S cyclin, Cyclin E, is required for CySC self-renewal; however, its canonical transcriptional regulator, a complex of the E2f1 and Dp transcription factors is dispensable for self-renewal and cell cycle progression. Nevertheless, we demonstrate that E2f1/Dp activity must be silenced to allow CySCs to differentiate. We show that E2f1/Dp activity inhibits the expression of genes important for mitochondrial activity. Furthermore, promoting mitochondrial activity or biogenesis is sufficient to rescue the differentiation of CySCs with ectopic E2f1/Dp activity but not their exit from the cell cycle. Our findings together indicate that E2f1/Dp coordinates cell cycle progression with stem cell identity by regulating the metabolic state of CySCs.

## Introduction

Adult stem cells maintain tissue homeostasis by balancing self-renewal and differentiation (Jones and Wagers, 2008). In most adult tissues, proliferative capacity is limited to self-renewing stem cells and progenitors or transit-amplifying cells, but terminally differentiated cells are post-mitotic (Ruijtenberg and van den Heuvel, 2016). How cell cycle state and cell identity are coordinated to achieve this distribution of proliferative capacity is still poorly understood.

During development, terminal differentiation is accompanied by permanent cell cycle exit, where cells maintain 2N DNA content and are refractory to entering S phase (Buttitta and Edgar, 2007; Ruijtenberg and van den Heuvel, 2016). Negative regulators of the G1-S transition function to inhibit proliferation in terminally differentiating cells as entry into S phase is the critical point after which cells commit to undergoing a round of division (Pardee, 1974; Wharton, 1983). During normal cycling, growth factor signalling activates the Cyclin D (CycD) - cyclin-dependent kinase 4 (Cdk4) complex which phosphorylates the Retinoblastoma (Rb) protein to inhibit its function (Fig. 1A) (Brehm et al., 1998; Kato et al., 1993; Matsushime et al., 1994). Rb normally binds a transcription factor, composed of a dimer of transcriptional activator E2f proteins with Dimerisation Partner (Dp), and represses transcription. When this inhibition is relieved, E2f/Dp drive transcription of S-phase genes, and the late G1 cyclin, Cyclin E (CycE) (Brehm et al., 1998; DeGregori et al., 1995; Ohtsubo et al., 1995). Together with its partner, Cdk2, CycE promotes S phase entry. Terminal differentiation is associated with transcriptional induction of Rb and cyclin-dependent kinase inhibitors (CKIs) which block S phase entry, often in multiply redundant ways (Buttitta and Edgar, 2007; Ruijtenberg and van den Heuvel, 2016). However, the links between differentiation and cell cycle are tissue-specific as in some instances, affecting cell cycle exit influences differentiation, whereas in others, it results in differentiated cells that bear all the gene expression and functional characteristics of differentiation that continue to proliferate. In part, these differences have been attributed to direct binding between cell cycle inhibitors such as Rb and tissue-specific transcription factors required for differentiation, resulting in a link between cell identity and cell cycle exit (Buttitta and Edgar, 2007; Ruijtenberg and van den Heuvel, 2016).

**Figure 1.**
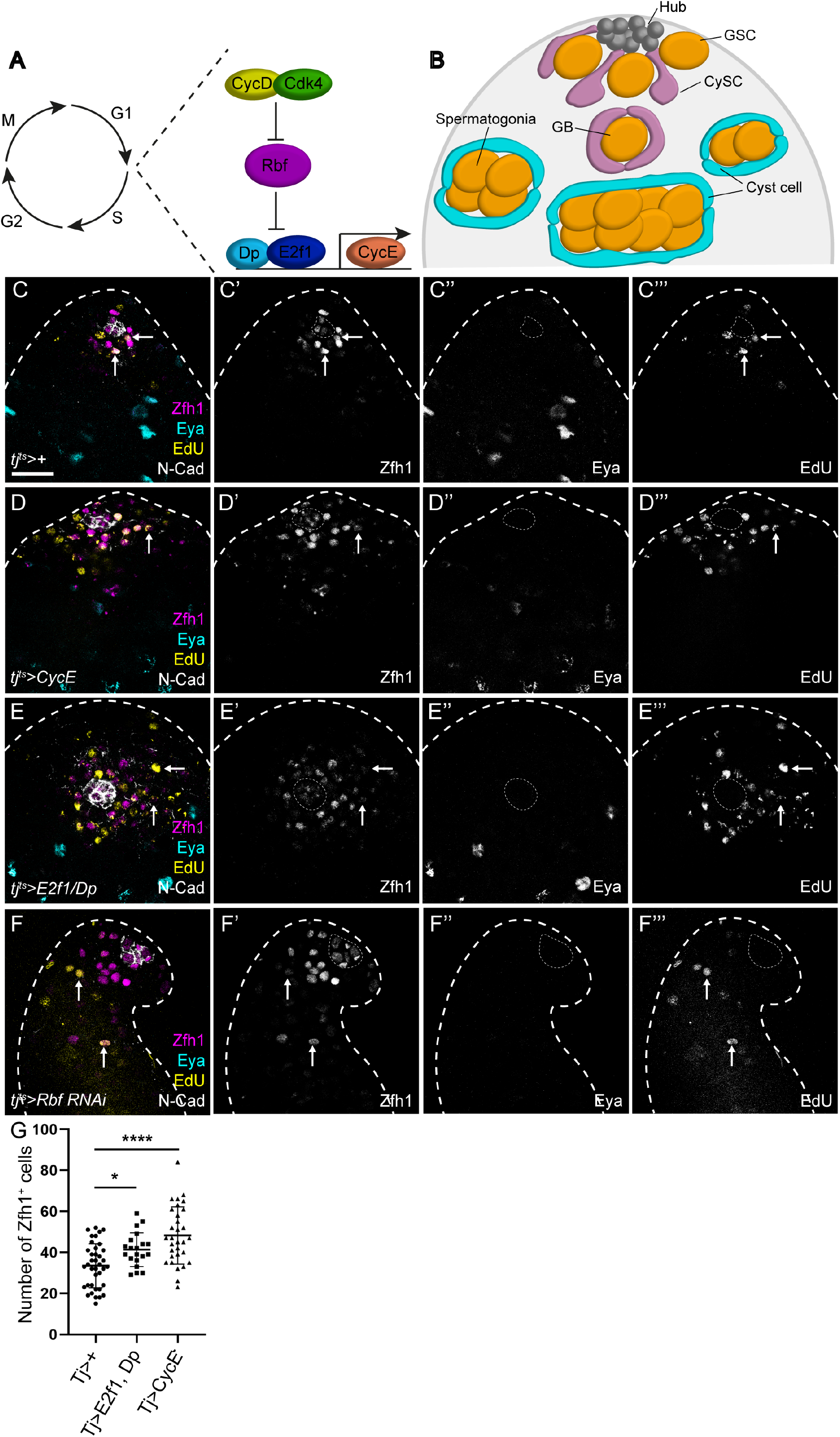
Promoting G1/S progression expands the CySC population. (A) Diagram of regulatory interactions controlling S phase entry. Cyclin D is expressed during G1 and together with Cdk4 phosphorylates Rbf to inactivate it. Rbf binds the E2f1/Dp dimer and represses transcription. Active E2f1/Dp promotes transcription of *Cyclin E*, which further inhibits Rbf and promotes entry into S phase. (B) Schematic of the Drosophila testis. The hub (grey) supports two stem cell population, both of which are identified by their contact with hub cells. Germline stem cells (GSCs, yellow) divide to give rise to gonialblasts that differentiate and undergo 4 rounds of incomplete division to produce 16 interconnected spermatogonia that will mature into spermatocytes. Somatic cyst stem cells (CySCs) give rise to cyst cells which wrap the spermatogonia and support their development. Cyst cells are post-mitotic, such that CySCs are the only proliferative somatic cells in the testis. (C) A control testis expressing *tj-Gal4* and *tub-Gal80*^*ts*^ (referred to as *tj*^*ts*^*>*) and labelled with antibodies against Zfh1 (magenta, single channel C’) to mark CySCs and early daughter cells, N-Cad (white) to label the hub, Eya (cyan, single channel C”) to label differentiated cyst cells, and EdU (yellow, single channel C’”), to mark cells in S phase. CySCs adjacent to the hub undergo S phase (arrows) while differentiated cyst cells never incorporate EdU. (D-F) Misexpression of CycE, (D) E2f1/Dp or (E) knockdown of Rbf (F) with *tj*^*ts*^*>* results in an increase of the number of Zfh1-expressing cells (magenta, single channels D’-F’) which are detected further from the hub. These cells incorporated EdU (yellow, single channels D’”-F’”) away from the hub (arrows), indicating ectopic proliferation. (G) Quantification of the number of Zfh1-expressing, Eya-negative cells. E2f1/Dp and CycE over-expression resulted in a significant increase in CySC numbers compared to control. * denotes P<0.05, **** denotes P<0.0001, determined with a Kruskal-Wallis test and Dunn’s multiple comparisons. Dotted lines outline the hub. Scale bar: 20µm in all panels.

In adult tissues, the links between proliferation and cell identity are similarly context-dependent. Many adult stem cells such as haematopoietic stem cells are quiescent; in these, inducing ectopic proliferation results in loss of self-renewal capacity (Cheng et al., 2000). Conversely, in highly proliferative stem cells such as those residing in Drosophila ovaries and testes, mutation of Cyclins, Cdks and the Cdk activator Cdc25 required for cell cycle progression results in loss of stem cell maintenance (Ables and Drummond-Barbosa, 2013; Inaba et al., 2011; Wang and Lin, 2005; Wang and Kalderon, 2009), suggesting that cell cycle progression promotes stem cell maintenance. Consistently, loss of the cell cycle inhibitor Rb results in expansion of the stem cell and progenitor populations and a block in terminal differentiation (Sage, 2012). Whether these functions of cell cycle regulators in self-renewal and differentiation are related to their roles in promoting cell cycle progression, or, as suggested for CycE (Ables and Drummond-Barbosa, 2013), whether these factors control identity and cell cycle progression independently is still unclear.

To gain insight into the mechanisms linking cell proliferation and identity in stem cells, we use the Drosophila testis as a model. The testis stem cell niche consists of a cluster of quiescent somatic cells called the hub, which is anchored to the apical tip of the testis and supports two stem cell populations. Germline stem cells (GSCs) divide to give rise to gonialblasts, which undergo a series of incomplete divisions to form a 16-cell cyst which matures and undergoes meiosis to form spermatids. A population of somatic stem cells, called cyst stem cells or CySCs, give rise to cyst cells which ensheath the developing germline and support its differentiation (Hardy et al., 1979; Kiger et al., 2000; Tran et al., 2000). CySCs are the only proliferating somatic cells; their daughters exit the cell cycle as they differentiate (Cheng et al., 2011; Gonczy and DiNardo, 1996; Hardy et al., 1979) (Fig. 1B), providing an ideal system to study how cell identity is linked to proliferative capacity. Previous work has shown a link between cell cycle progression and CySC identity: CySCs lacking the Cdk activator *cdc25* (called *String* in Drosophila) are not maintained (Inaba et al., 2011), while conversely, accelerating proliferation results in an increased likelihood of self-renewal at the expense of neighbouring wild type CySCs (Albert et al., 2018; Amoyel et al., 2016; Amoyel et al., 2014; Michel et al., 2012). Moreover, mutants for the Rb homologue *Rbf* accumulate stem-like cells and lack differentiated cells in larval testes (Dominado et al., 2016). In adults, *Rbf* is required to maintain quiescence of terminally differentiated hub and cyst cells (Greenspan and Matunis, 2018), although its role in adult stem cells has not been established.

Since differentiating daughters of CySCs exit the cell cycle without transit-amplifying divisions, we asked whether positive regulators of G1-S transition were important in maintaining CySC identity and, conversely, whether inhibitors of G1-S progression were required to coordinate cyst cell differentiation with cell cycle exit. We took advantage of reduced genetic redundancy in Drosophila, which has one activator E2f, called E2f1, and one repressor E2f, E2f2, both of which bind a single DP homologue, Dp, which mediates transcription by both activator and repressor complexes (Dynlacht et al., 1994; Frolov et al., 2001; Frolov et al., 2005; Korenjak et al., 2012; Sawado et al., 1998).

We find that CycE is necessary for CySC self-renewal and its over-expression leads to increased CySC numbers and proliferation. Similarly, loss-of-function of *Rbf* results in ectopic CySC-like cells throughout the testis in an E2f1/Dp-dependent manner. However, E2f1 and Dp, which in other contexts are required to promote *CycE* expression, are dispensable for normal CySC cycling, despite being endogenously present and active, implying that silencing of E2f1/Dp by Rbf is essential to allow CySCs to differentiate. We compare the transcriptional changes induced by Rbf depletion to those occurring in normal differentiation and show that differentiation is accompanied by changes in metabolic gene expression. Finally, we show that differentiation, but not ectopic proliferation of Rbf-deficient cells can be rescued by co-expression of transcription factors that promote mitochondrial biogenesis, thereby uncoupling differentiation and cell cycle progression. Altogether, we conclude that cell cycle progression is essential for CySC maintenance, but that the E2f1/Dp complex is not involved in normal cycling or CySC maintenance. We therefore propose that E2f1/Dp activity in CySCs coordinates cell cycle progression with stem cell identity by controlling CySC metabolism.

## Results

### Promoting the G1/S transition causes ectopic proliferation and expands the CySC population

To test whether entry into S-phase was linked to maintenance of CySC identity, we asked whether promoting progress through the G1/S transition could also affect cell fate. To this end, we manipulated the key Cyclin controlling S phase entry, CycE, and the transcriptional regulator of S-phase genes, the E2f activator complex, composed of E2f1 and Dp and their inhibitor, Rbf. We assessed cell proliferation by EdU incorporation and cell identity using antibodies against Zfh1, which labels CySCs and their immediate daughters (Leatherman and Dinardo, 2008), and Eya, which labels post-mitotic, differentiated cyst cells (Fabrizio et al., 2003). In control testes, Zfh1 expression was detected in two tiers of cells surrounding the hub, whereas cyst cells distant from the hub expressed Eya. Consistent with prior reports that the only proliferating somatic cells in the testis are CySCs (Cheng et al., 2011; Gonczy and DiNardo, 1996; Hardy et al., 1979), DNA replication in somatic cells, as assayed by EdU incorporation, was only detected in Zfh1-positive cells around the hub (Fig. 1C, arrows).

We mis-expressed CycE in the cyst lineage using *traffic jam* (*tj*)*-Gal4*, together with a ubiquitously-expressed temperature-sensitive Gal80 (referred to as *tj*^*ts*^). Restricting mis-expression to adult flies by growing embryos and larvae at the permissive temperature and shifting them as adults to the restrictive temperature of 29°C for 10 days resulted in somatic cells distant from the hub incorporating EdU (Fig. 1D, arrows), consistent with a role for CycE in promoting S-phase entry. In addition, we observed an expansion of Zfh1-expressing cells away from the niche compared to controls (Fig. 1D). The number of Zfh1-positive cells increased significantly in CycE-expressing testes compared to controls (Kruskal-Wallis followed by Dunn’s multiple comparison test, P<0.0001, Fig. 1G), from 33.5±1.7 (n=38 testes) to 48.3±2.5 (n=31). However, we always observed Eya-positive cells further distally (Fig. 1D), and never observed EdU incorporation in these cyst cells, suggesting that, although delayed, differentiation occurred normally in somatic cells over-expressing CycE.

Next, we tested whether the transcriptional regulator of S-phase gene expression, E2f1, together with its partner Dp could influence CySC fate. Previous work has suggested that E2f1/Dp is active in CySCs (Amoyel et al., 2014). Using an established reporter for E2f1/Dp transcriptional activity, *PCNA-GFP* (Thacker et al., 2003), we detected E2f1/Dp activity in CySCs (Fig. S1A, arrows), but not in differentiated cyst cells away from the hub (Fig. S1A, arrowheads). In CySCs, *PCNA-GFP* was detected in a subset of cells, consistent with periodic cell cycle-dependent activation of E2f1/Dp. Reporter expression was strongly reduced upon Dp knockdown (Fig. S1B), indicating that *PCNA-GFP* expression reflects endogenous Dp-dependent transcription. Similar to CycE over-expression, Dp and E2f1 over-expression led to ectopic proliferation of somatic cells far from the hub (Fig. 1E, arrows). In addition, we counted 41.4±1.8 Zfh1-positive cells, significantly higher than the control (n=20, Kruskal-Wallis followed by Dunn’s multiple comparison test, P<0.046, Fig. 1G). To test whether activating endogenous E2f1/Dp could also result in ectopic Zfh1-expressing cells, we knocked down the negative regulator of this complex, Rbf. Rbf expression was detected in all cells at the apical tip of the testes (Fig S2A), as previously described (Greenspan and Matunis, 2018), and efficient knockdown was achieved by RNAi expression (Fig S2B). Previous work has shown a role for *Rbf* in the cyst lineage during development (Dominado et al., 2016), where loss of Rbf led to ectopic proliferation and a lack of differentiated cyst cells, while in adult testes, Rbf was reported to maintain quiescence of differentiated cyst cells (Greenspan and Matunis, 2018). Expression of an RNAi against *Rbf* in the somatic lineage led to an expansion of the Zfh1-positive population, in addition to ectopic proliferation away from the hub (Fig. 1F). Importantly, testes in which Rbf was knocked down had no Zfh1-negative, Eya-positive cyst cells (Fig. 1F), implying a complete block in differentiation.

Altogether, over-activating key promoters of the G1/S transition is sufficient to promote proliferation away from the stem cell niche and affects the ability of CySCs to differentiate.

### *CycE* is required for CySC self-renewal

Next, we asked whether these regulators were necessary for the maintenance of CySC identity. We generated loss-of-function CySC clones using mitotic recombination. Since CySCs are the only dividing somatic cells in the testis, any labelled somatic cells were necessarily generated from a CySC division. We measured the persistence of clones over time as a reflection of the ability of labelled CysCs to self-renew in the niche. Clonal CySCs were identified as GFP-positive, Zfh1-positive cells adjacent to the hub. Control marked clones were readily recovered at 2 days post clone induction (dpci) (Figs. 2A, 2F, Table 1) and were maintained at 7 dpci (Figs. 2B, 2F, Table 1).

**Table 1.** Clone recovery rates (% testes) for the indicated genotypes. Clone recovery rates that are significantly different from their respective controls are indicated by blue shading. All others are not significantly different, as determined by Fisher’s exact test. N.B. the *Cdk4*^*s4639*^ experiment was carried out separately from other FRT^42D^ experiments with an independent control. ND: Not determined.

| Genotype | 2 dpci<br>(n) | 7 dpci<br>(n) | 14 dpci<br>(n) |
| --- | --- | --- | --- |
| Control (*FRT*<sup>40A</sup>) | 43<br>(23) | 48<br>(54) | ND |
| *CycE*<sup>AR95</sup> | 16<br>(31) | 0<br>(37) | ND |
| Control (*FRT*<sup>42D</sup>) | 67<br>(95) | 55<br>(148) | 65<br>(46) |
| *Dp*<sup>a3</sup> | 67<br>(18) | 56<br>(55) | 66<br>(29) |
| *Dp*<sup>a4</sup> | ND | 62<br>(47) | 67<br>(30) |
| *Cdk4*<sup>k06503</sup> | 53<br>(38) | 47<br>(34) | ND |
| Control (*FRT*<sup>42D</sup>) | 67<br>(42) | 50<br>(42) | ND |
| *Cdk4*<sup>s4639</sup> | ND | 33<br>(52) | ND |
| Control (*FRT*<sup>82B</sup>) | 42<br>(45) | 45<br>(93) | ND |
| *E2f1*<sup>rM729</sup> | 45<br>(33) | 33<br>(55) | ND |

**Figure 2.**
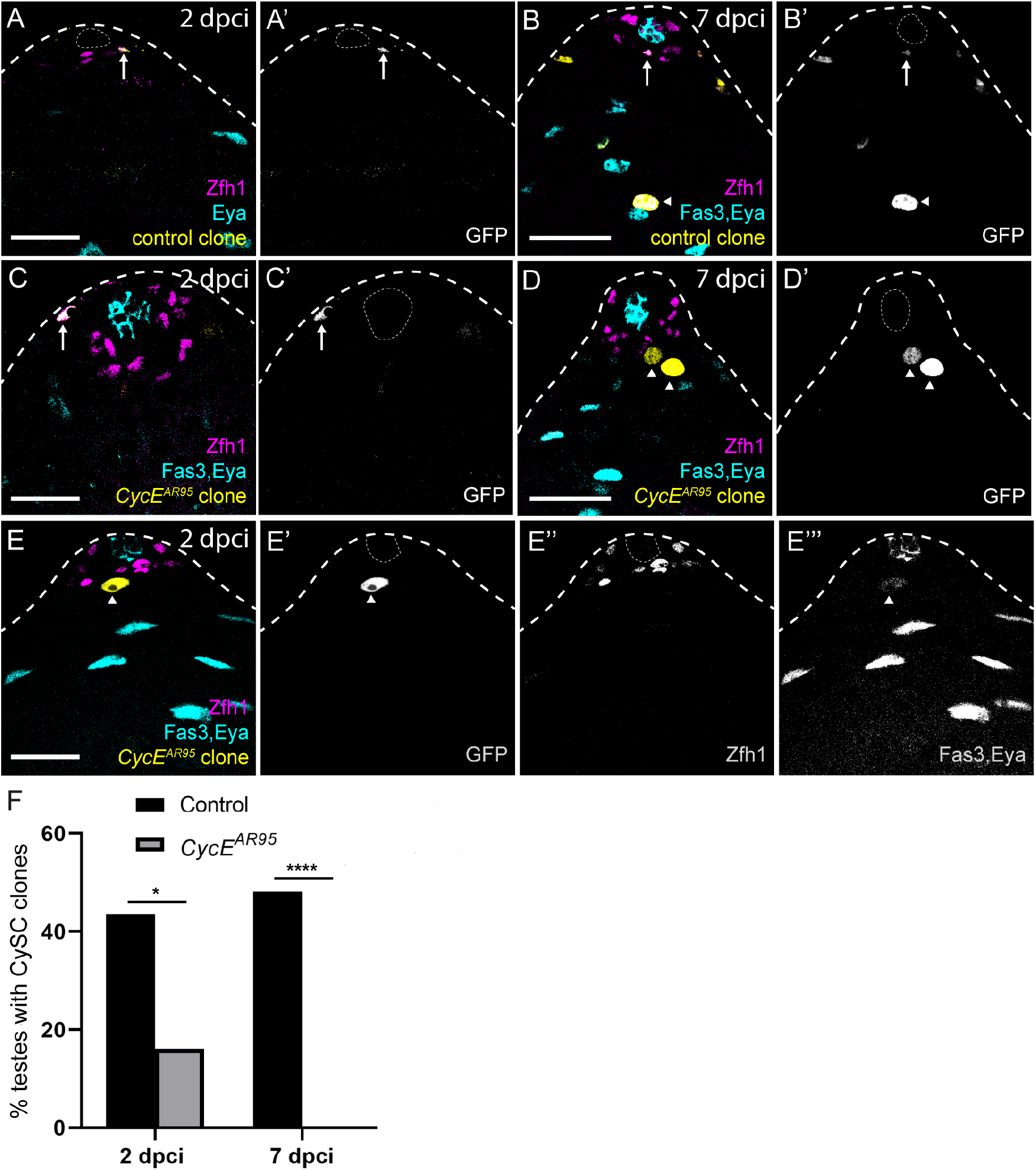
*CycE* mutant clones differentiate prematurely. (A-E) Control (A,B) or *CycE* (C-E) mutant clones were positively labelled with GFP (yellow, single channel in A’-E’) and identified as CySCs by Zfh1 expression (magenta) or cyst cells using Eya (cyan). (A,B) control clones were recovered at 2 days post clone induction (dpci,) and maintained at 7 dpci (B). (C,D) *CycE*^*AR95*^ mutant CySCs were observed at 2 dpci (arrow) but by 7 dpci no *CycE*^*AR95*^ mutant clones expressed Zfh1 and only Eya-expressing cells were detected (arrowheads). Note the enlarged nucleus of mutant cells. (E) *CycE*^*AR95*^ mutant cells at 2 dpci expressed Eya (single channel E’”) prematurely and downregulated Zfh1 (single channel E”), while neighbouring cells at a similar distance from the hub maintained Zfh1 expression. (F) Graph showing the fraction of testes containing marked control or *CycE*^*AR95*^ mutant CySC clones. See Table 1 for n values. * denotes P<0.05, **** denotes P<0.0001, determined by Fisher’s exact test. Dotted lines outline the hub. Scale bars: 20µm.

By contrast, CySC clones mutant for a null allele of *CycE* were never recovered at 7 dpci (Figs. 2C-D, 2F, Table 1). The impaired self-renewal of *CycE* mutant CySCs was already evident at 2 dpci, as few CySC clones were observed at that stage (P<0.035, Fisher’s exact test, Fig. 2F, Table 1). By 7 dpci, only Eya-positive differentiated cells were labelled (P<0.0001, Fisher’s exact test, Fig. 2D, arrowheads). We note that Eya-positive *CycE* mutant cells did not appear wild type, as their nuclei were enlarged, similar to reports in the female germline (Ables and Drummond-Barbosa, 2013). Despite low CySC clone recovery rates even at 2 dpci, clones were induced at similar rates to controls, as GFP-positive *CycE* mutant somatic cells were observed in 26/31 (or 84%) testes examined compared to 20/23 (87%) in controls (P>0.99, Fisher’s exact test). However, most of these cells did not express Zfh1, indicating that *CycE* mutant CySCs differentiated rapidly. We sought to establish the cause of the low recovery of *CycE* mutant CySCs. By 2 dpci, we observed mutant clones that expressed Eya prematurely (Fig. 2E, arrowhead), surrounded by wild type cells expressing Zfh1. This observation suggests that *CycE* mutant CySC clones are poorly recovered because they differentiate prematurely.

Taken together with the gain-of-function experiments (Fig. 1), our data indicate that *CycE* is necessary for CySC self-renewal and at least partly sufficient to drive ectopic self-renewal, establishing *CycE* as a critical regulator of CySC fate.

### The E2f1/Dp complex is not required for CySC self-renewal

In most contexts studied to date, *CycE* expression is transcriptionally induced at the G1/S transition by the E2f1/Dp complex (Dimova and Dyson, 2005; Dimova et al., 2003; Duronio et al., 1996; Duronio et al., 1995; Korenjak et al., 2012; Stevaux and Dyson, 2002; Vermeulen et al., 2003). Since *CycE* is essential for CySC maintenance, we reasoned that *E2f1* and *Dp* would also be required for CySC self-renewal and generated mutant clones to assess their role.

Unexpectedly, both control and *Dp* mutant CySC clones were recovered at 7 and 14 dpci (Fig. 3B,C) with indistinguishable clone recovery rates (P>0.99 at both 7 and 14 dpci for *Dp*^*a3*^ and P=0.72 at 7 dpci and P<0.99 at 14 dpci for *Dp*^*a4*^ compared to control, Fisher’s exact test, Fig. 3A-D). We confirmed this surprising result with two separate null alleles (Fig. 3D, Table 1), and verified genetically that the alleles we used were indeed *Dp* mutants (See Materials and Methods). Moreover, we examined the expression of the Ef21/Dp transcriptional target *PCNA* in *Dp* mutant clones and observed reduced reporter expression (Fig. 3E). Dp/E2f1 transcription is thought to drive S-phase progression, yet the presence of *Dp* mutant clones at 7 and 14 dpci encompassing many Zfh1-positive CySCs and Eya-positive cyst cells implied that these clones had proliferated. We confirmed this directly by observing EdU incorporation in *Dp* mutant clones both at 7 and 14 dpci (Fig. 3F,G), indicating that *Dp* is dispensable for DNA replication in CySCs. Although surprising, this result is consistent with work showing that most tissues in *Drosophila* can proliferate in the absence of Dp activity during development (Frolov et al., 2001; Royzman et al., 1997; Zappia and Frolov, 2016).

**Figure 3.**
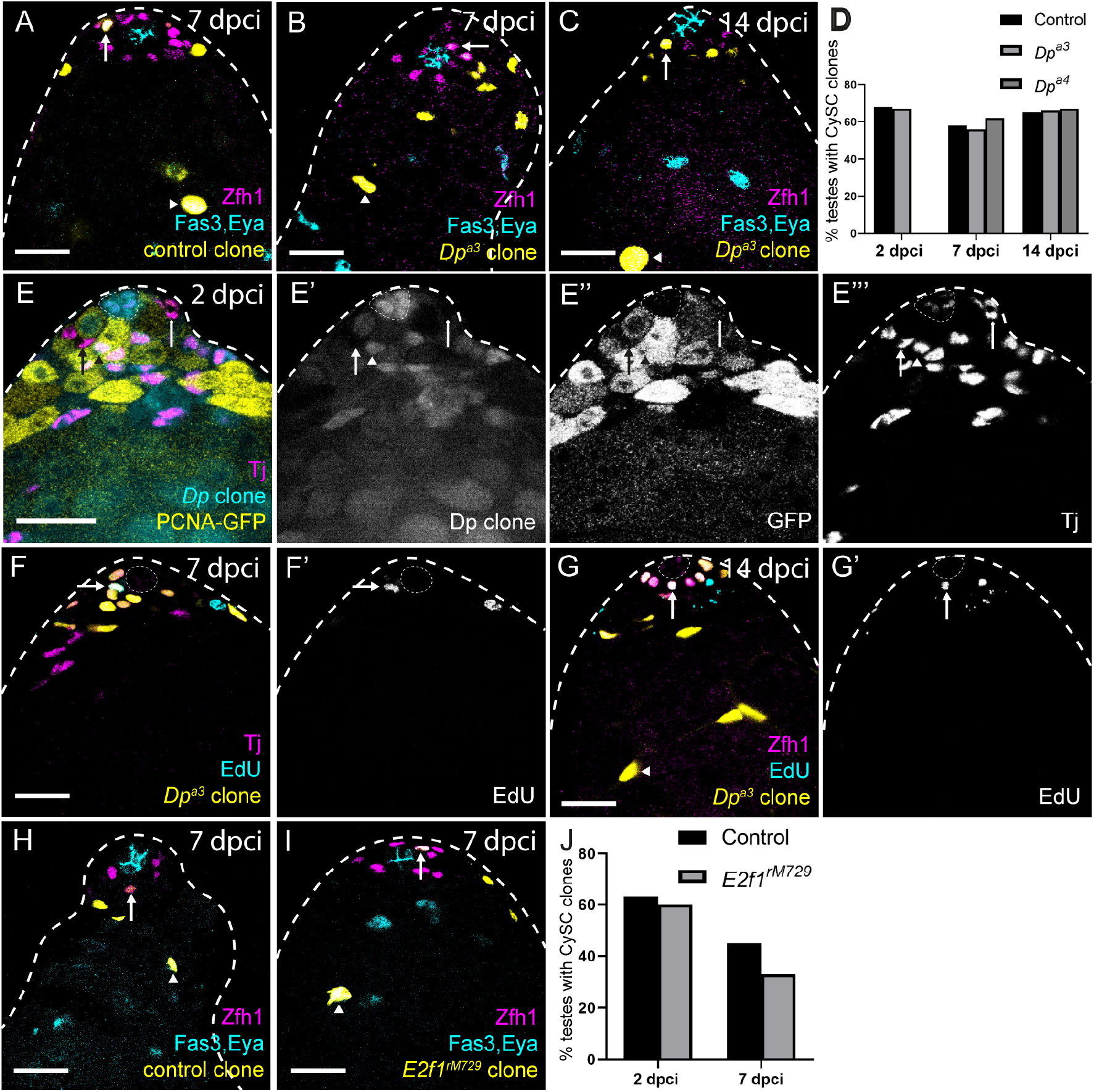
Dp and E2f1 are dispensable for CySC self-renewal. Positively-marked control (A,H), *Dp* mutant (B,C,F,G) or *E2f1* mutant (I) CySCs labelled with GFP expression (yellow) at the indicated days post clone induction (dpci). CySCs were identified by Zfh1 expression (magenta) and position adjacent to the hub (Fas3, cyan). Differentiated cyst cells were labelled with Eya (cyan). (A-D) Control clones (A) and *Dp* mutant (B,C) clones were recovered at similar rates, and maintained over time. (D) Graph showing the fraction of testes containing marked control or *Dp* mutant CySC clones. Clone recovery rates were not significant different at 7 and 14 dpci, determined by Fisher’s exact test. See Table 1 for n values. (E) Negatively-marked *Dp* mutant clones at 2 dpci labelled by lack of RFP expression (cyan, single channel in E’). Mutant CySCs (arrows) displayed decreased levels of PCNA-GFP expression (yellow, single channel in E”), a canonical transcriptional target of Dp. Somatic cells including CySCs are labelled with Tj (magenta, single channel in E’”). (F,G) Positively-labelled *Dp* mutant clones at 7 dpci (F) and 14 dpci incorporated EdU, indicating that they could enter S phase. (H-J) *E2f1* mutant clones (I) were recovered at 7 dpci, similar to controls (H). (J) Quantification of clone recovery rates, showing the fraction of testes containing control or *E2f1*^*rM729*^ CySC clones. Mutant clone recovery rates were not significant different from controls, determined by Fisher’s exact test. See Table 1 for n values. Dotted lines outline the hub. Scale bars: 20µm.

Consistently, *E2f1* mutant CySC clones were recovered at 7dpci (Fig. 3H-I, arrows) with a slightly lower but not significantly different clone recovery rate to controls (Fisher’s exact test, P>0.05, Fig. 3J). Like control and *Dp* mutant clones, *E2f1* mutant clones were composed of both Zfh1-expressing CySCs and Eya-positive differentiating cyst cells (Fig. 3H,I, arrowheads), suggesting that neither their self-renewal nor their differentiation capacity was altered.

Finally, we tested whether an upstream positive regulator of E2f1/Dp was required for CySC self-renewal. CycD/Cdk4 activity induces E2f1/Dp activity by phosphorylating and inhibiting Rbf. We induced mutant clones for two separate alleles of *Cdk4*. These clones were recovered at similar rates to controls at 7 dpci (Fig. S3, Table 1), indicating that *Cdk4* is not required for CySC self-renewal.

Altogether, our data show that E2f1/Dp activity is not required in CySCs for self-renewal, despite the fact that ectopically activating this complex is sufficient to drive CySC over-proliferation (Fig. 1). This result stands in contrast with the requirement identified above for *CycE* in CySC self-renewal and implies that, contrary to the prevalent model of cell cycle regulation, *CycE* expression in CySCs does not depend on E2f1/Dp or CycD/Cdk4 activity. Instead, our data suggest that *CycE* is either constitutively expressed or that other transcriptional regulators take the role usually assigned to E2f1/Dp.

### E2f1/Dp inhibition by Rbf is required for cyst cell differentiation

Despite a genetic lack of requirement for *Dp* in CySC self-renewal, Dp-dependent transcriptional activity was detected in CySCs (Fig. S1) (Amoyel et al., 2014), in a pattern suggesting cell cycle-dependent E2f1/Dp activation. Moreover, Rbf knockdown resulted in a block of CySC differentiation (Fig. 1F), suggesting that E2f1 and Dp were indeed expressed in the cyst lineage. To resolve the question of what endogenous role Rbf and E2f1/Dp activity might play in CySC self-renewal and differentiation, we characterised the role of *Rbf* in CySCs and examined the dependency of Rbf knockdown on E2f1/Dp.

First, we characterised the phenotype of *Rbf* loss-of-function in CySCs, both in clonal and lineage-wide knockdown experiments. Lineage-wide Rbf knockdown with *tj*^*ts*^ led to expansion of Zfh1-expressing cells such that they filled the entire testis, and an absence of Eya-positive differentiated cyst cells (Fig. 1F, Figs. 4A-B). Similarly, CySC clones over-expressing two different RNAi constructs against Rbf were entirely composed of Zfh1-expressing cells and devoid of Eya-expressing cells (Fig. S4A-C). Although *Rbf* is X-linked, we generated negatively-marked CySC clones hemizygous mutant for the loss-of-function allele *Rbf*^*14*^, using a rescuing duplication on an autosome (see Methods). CySC clones mutant for *Rbf* displayed a similar phenotype to Rbf-knockdown CySCs: clones were only composed of Zfh1-expressing cells and no Eya-expressing differentiated cyst cells were present within the clones (Fig. S4D,E, n=22/26), indicating a block in differentiation in *Rbf* mutant CySCs.

**Figure 4.**
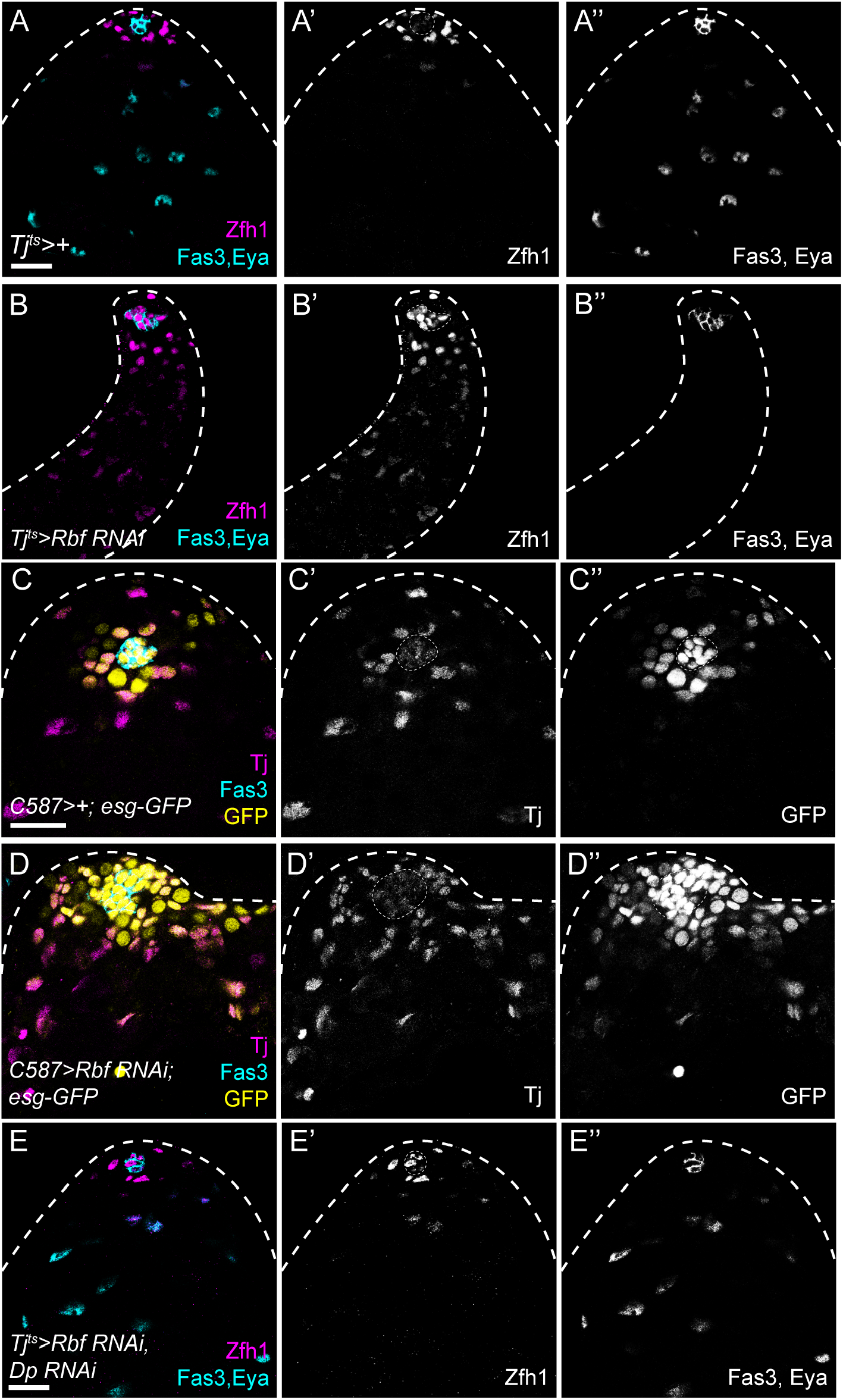
Inhibition of E2f1/Dp is necessary for cyst cell differentiation. (A) Control testis showing Zfh1 expression (magenta, single channel A’) in CySCs surrounding the hub (labelled with Fas3, cyan, single channel in A”). As cells differentiate, Zfh1 expression is downregulated and cyst cells express Eya (cyan, single channel in A”). (B) Knockdown of Rbf with *tj*^*ts*^ results in ectopic Zfh1 expression away from the hub (magenta, single channel in B’) and a lack of Eya expression (cyan, single channel in B”). (C,D) *esg-GFP* expression (yellow, single channels in C”,D”) in control (C) and Rbf knockdown testes (D). The hub is labelled with Fas3 (cyan), and CySCs and early cyst cells are labelled with Tj (magenta, single channels in C’,D’). In controls (C) GFP is expressed in the hub, GSCs and occasional early differentiating gonia, and in somatic CySCs adjacent to the hub. Expression of *esg-GFP* is detected in somatic cells distant from the hub in Rbf knockdowns (D). (E) Knockdown of both Dp and Rbf resulted in Zfh1 expression (magenta, single channel in E’) being restricted around the hub (Fas3, cyan) and Eya-expressing cells detected further from the hub (cyan, single channel in E”), similar to control testes, and indicating that Dp loss suppresses the differentiation block induced by Rbf knockdown. Dotted lines outline the hub. Scale bars: 20µm.

The ectopic Zfh1-expressing, Eya-negative cells that were detected away from the niche in Rbf knockdowns expressed low levels of Zfh1 compared to cells adjacent to the hub (Figs. 1F, 4B). To confirm whether these ectopic cells were indeed CySCs, we examined other markers of CySCs, Esg, Chinmo and Wg (Flaherty et al., 2010; Leatherman and Dinardo, 2008; Voog et al., 2014). In controls, both Esg and Chinmo labelled CySCs, as well as the hub and early germ cells. Both markers were expanded throughout Rbf-knockdown testes and showed an expression pattern similar to Zfh1 (Figs. 4C-D, S5A,B), with high expression in CySCs close to the hub and lower levels in ectopic cells throughout the testis. Wg expression was observed in distinctive puncta around the hub in control testes, but punctate expression was observed throughout the testis in Rbf knockdowns (Fig. S5C,D). These results suggest that Rbf-depleted cells maintain a CySC-like state, similar to immediate CySC daughters which have lower Zfh1 expression than CySCs in contact with the hub (Leatherman and Dinardo, 2008), but that their differentiation does not progress further. Furthermore, we ruled out cell death and ectopic self-renewal signalling as explanations for the role of Rbf in promoting CySC differentiation (see Supplementary text and Figs S6 and S7).

Previous work has shown that, in larvae, Rbf binding to DNA is abolished in the absence of Dp (Korenjak et al., 2012), indicating that Rbf exerts its effects on gene expression through Dp. Indeed, loss-of-function of both *Dp* and *Rbf* led to a similar distribution of Zfh1-expressing cells around the hub and Eya-expressing cells away from the hub as in control testes, both in tissue-wide knockdowns (Fig. 4E) and in mutant clones (Figs. S4G, S4H), despite lacking any detectable Rbf protein by immunohistochemistry (Fig. S4I, arrowhead). Consistently, *E2f1* loss-of-function also rescued the lack of differentiation of Rbf RNAi clones (Figs. S4F, S4H), similar to observations in larval testes (Dominado et al., 2016). Thus, the effects of *Rbf* loss-of-function in CySCs are attributable entirely to ectopic E2f1/Dp activity.

In sum, our data suggest that the E2f1/Dp complex acts as a link to coordinate cell cycle and CySC identity. While E2f1/Dp activity is not required for cell cycle progression, it inhibits differentiation in CySCs progressing through S phase. In turn, Rbf acts as a permissive factor for differentiation by relieving the E2f1/Dp-mediated inhibition of differentiation.

### Ectopic E2f1/Dp activity in CySCs alters expression of genes regulating metabolism and energy production

To gain insight into the mechanisms by which the E2f1/Dp complex transcriptionally inhibits differentiation, we compared gene expression in sorted somatic cells in *tj>+* control and Rbf knockdown testes. Using stringent criteria (2-fold change and FDR<0.01), we identified >5000 differentially-expressed transcripts (3329 upregulated and 2098 downregulated in Rbf knockdown compared to control) (Fig. 5A, Supplementary File S1), indicating that Rbf knockdown results in a large disruption to gene expression, presumably as a combination of the direct effects of Rbf loss and indirect effects of a block in differentiation.

**Figure 5.**
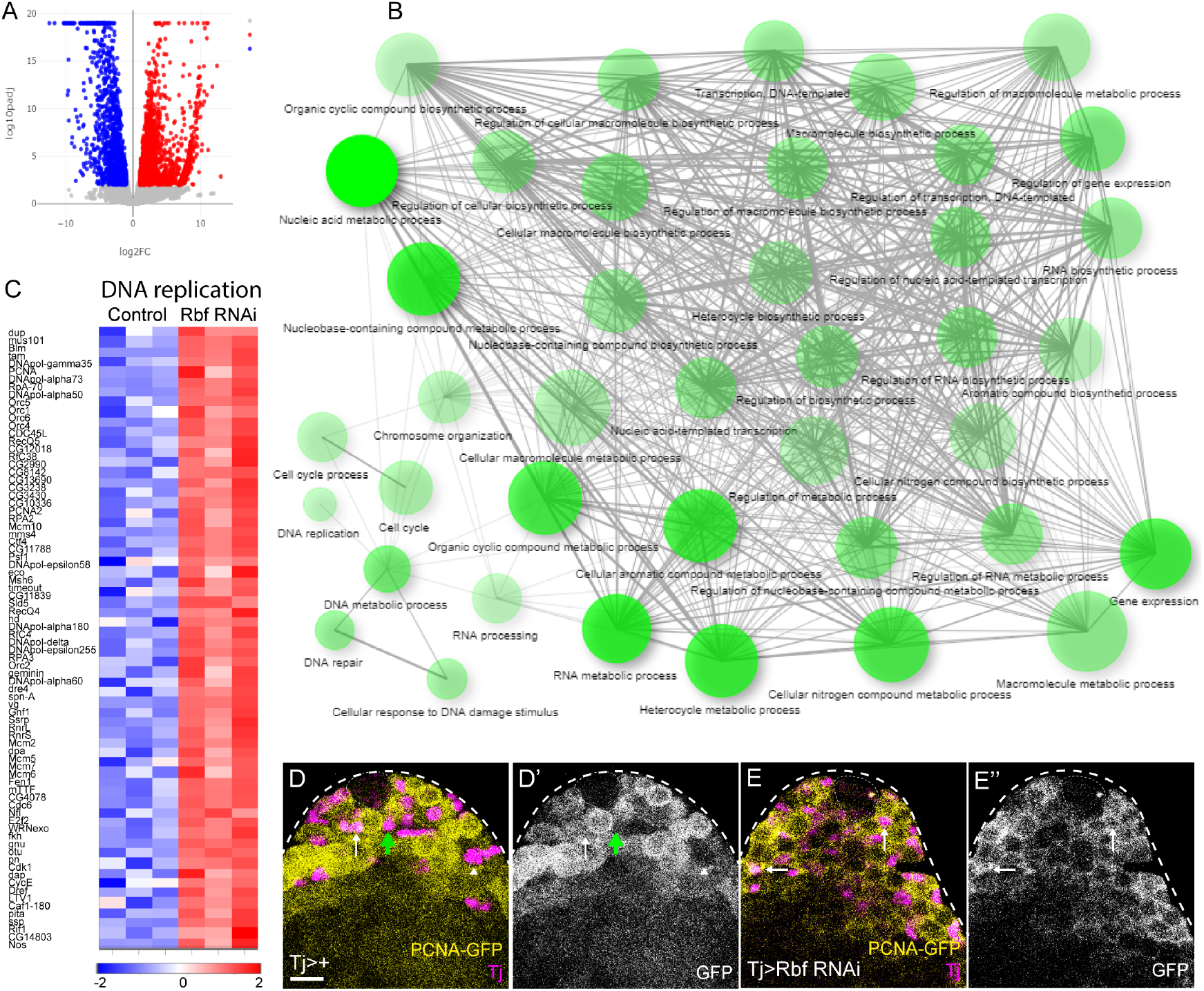
Rbf knockdown results in upregulation of genes involved in cell cycle progression and DNA replication. (A) Volcano plot showing log_10_ of the adjusted P value (log_10_(p_adj_)) against the log_2_(fold change) for individual genes when comparing expression from sorted somatic cells from Rbf knockdown to control testes. Genes whose expression was 2-fold downregulated and upregulated with a false discovery rate (FDR) below 0.01 are labelled in blue and red, respectively. (B) Gene ontology analysis of genes upregulated upon Rbf knockdown. The size of the circles represents the number of genes in each biological process. A higher significance of enrichment is shown as brighter colouring of the circle. Lines connect nodes with at least 15% of shared genes and their thickness increases with the percentage of shared genes. This analysis reveals that Rbf knockdown results in upregulation of many genes involved in categories related to DNA metabolic processes, nucleotide synthesis and transcription. (C) Heat map showing relative expression of genes grouped under the GO term “DNA replication” for control and Rbf knockdown testes. Columns represent biological replicates. Colour key shows expression change in log_2_(fold change); blue represents lower expression, while higher expression is shown in red. (D,E) Expression of the *PCNA-GFP* reporter is increased in Rbf knockdown testes. In controls (D), GFP (yellow, single channel in D’) is detected in a subset of Tj-positive cells (magenta, white arrow) adjacent to the hub, and absent from differentiated cyst cells (white arrowhead). The black arrow indicates a Tj-positive CySC adjacent to the hub that does not express GFP. In Rbf knockdown testes (E) GFP expression is detected in Tj-positive cells (magenta) distant from the hub (white arrows). Dotted lines outline the hub. Scale bars: 20µm.

Gene ontology (GO) analysis of upregulated genes revealed an enrichment for several processes involved in cell cycle progression, DNA synthesis and replication (Fig 5B). The latter category included many well-described E2f1/Dp targets, such as *PCNA* and *Minichromosome maintenance* (Mcm) genes (Ishida et al., 2001; Yamaguchi et al., 1995) (Fig. 5C). In particular, *PCNA* was upregulated 20-fold following Rbf knockdown (Supplementary file S1), which we confirmed using the *PCNA-GFP* reporter (Figs. 5D-E). In controls, GFP in the somatic lineage was observed only adjacent to the hub (Fig. 5D, white arrow), whereas it was absent far from the hub (Fig. 5D, arrowhead). However, in Rbf knockdowns, GFP was detected in somatic cells distant from the hub (Fig. 5E, arrows). In addition to cell cycle-related genes, expression of CySC and early cyst cell markers was also differentially detected in the Rbf knockdown, including *Zfh1* (10.4 fold increase, FDR<0.001), *chinmo* (5.2 fold increase, FDR<0.001), and *tj* (17.9 fold increase, FDR<0.001). These results recapitulate our previous observations that these markers were ectopically expressed in Rbf knockdowns (Figs. 1D, S5) and suggest that our sequencing approach identified both likely direct Dp/E2f1 transcriptional targets and indirect targets that are upregulated as a consequence of an increase in the representation of CySC-like cells in the samples.

While upregulated transcripts were largely consistent with increased proliferation, transcripts that were downregulated upon Rbf knockdown fell into distinct categories (Fig. 6A). Transcripts encoding cell junction proteins were downregulated, in keeping with previous reports showing that expression of septate and adherens junction components increases during cyst cell differentiation (Dubey et al., 2019; Fairchild et al., 2017; Fairchild et al., 2015; Papagiannouli et al., 2019). A large number of downregulated transcripts encoded genes involved in various aspects of oxidative metabolism and ATP production (Fig. 6A,B), suggestive of an altered metabolic state in Rbf-deficient cells. To validate the downregulation observed by sequencing, we examined the expression of a GFP protein trap in the *Aldolase 1 (Ald1)* locus, which was downregulated in the Rbf knockdown group. In controls, Ald1-GFP expression was detected at low levels in CySCs and increased distally from the hub (Fig. 6C). Expression was reduced in both CySCs adjacent to the hub and ectopic CySC-like cells in testes in which Rbf was knocked down somatically (Fig. 6D).

**Figure 6.**
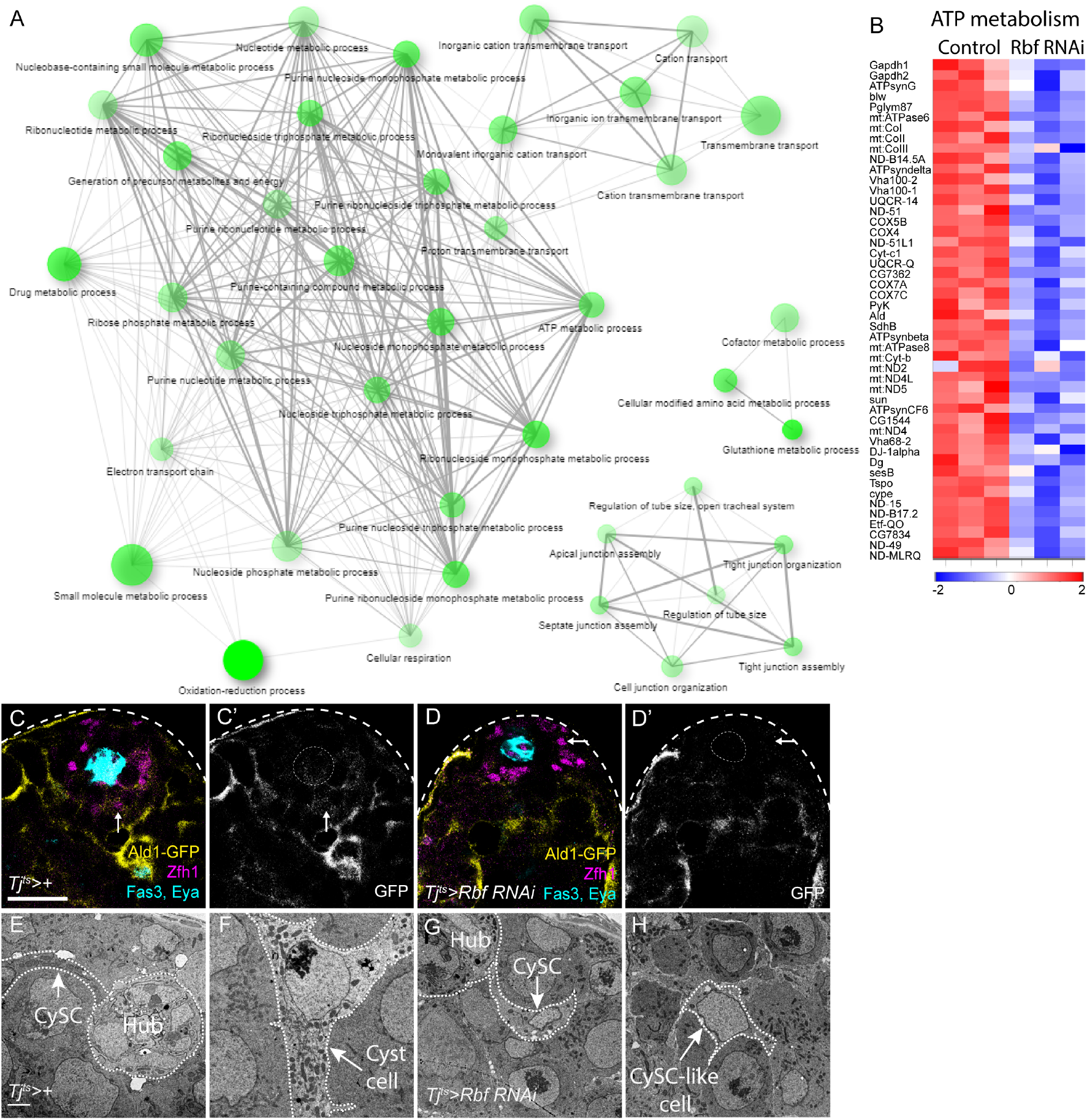
Rbf knockdown downregulates genes related to oxidative metabolism and energy production. (A) Gene ontology analysis of genes downregulated in Rbf knockdown testes. The size of the circles represents the number of genes in each biological process. A higher significance of enrichment is shown as stronger colouring of the circle. Lines connect nodes with at least 15% of shared genes and their thickness increases with the percentage of shared genes. Significant enrichment is observed for many categories related to oxidation-reduction and ATP metabolism. (B) Heat map showing relative expression of genes grouped under the GO term “ATP metabolism” for each biological replicate of control and Rbf knockdown sequencing (in columns Colour key shows expression change in log_2_(fold change); blue represents lower expression, while higher expression is shown in red. (C,D) Expression of a GFP protein fusion with the glycolytic enzyme Aldolase1 (Ald-GFP, yellow, single channels in C’,D’). In wild-type testes (C), GFP is detected at low levels in Zfh1-expressing CySCs (magenta, arrow), and at higher levels in differentiating cyst cells (labelled with Eya, cyan). In Rbf knockdowns (D), GFP expression is decreased and almost undetectable around the hub (Fas3, cyan). Scale bars in C,D: 20µm. (E-H) Electron micrographs of the testis apex in controls (E,F) and Rbf RNAi (G,H). A control CySC (E) can be seen contacting the hub (arrow). Many mitochondria are visible in the CySC cytoplasm along the length of its protrusion contacting the hub. In control differentiating cyst cells (F), numerous electron-dense, elongated mitochondria can be seen. In contrast, in an Rbf-deficient CySC (G), few mitochondria are seen in the cytoplasm and the projection contacting the hub. Similarly, in CySC-like cells distant from the hub in Rbf knockdowns (H), few mitochondria are present. Scale bars in E-H: 2µm.

Since reduced Ald1 expression in Rbf-deficient CySCs suggested that they may have a metabolic defect prior to differentiation, we examined mitochondrial morphology in control and Rbf-deficient CySCs using electron microscopy. In control testes, the cytoplasm of CySCs contained numerous mitochondria (Fig. 6E). Mitochondria in differentiated cyst cells were more elongated and more electron dense than in CySCs (Fig. 6F). By contrast, in Rbf knockdown CySCs, we observed many fewer mitochondria, and these appeared smaller and more globular than in controls (Fig. 6G). Somatic cells located away from the hub displayed similar small and round mitochondria and had many fewer mitochondria present than control differentiated cyst cells, resembling more closely the distribution and morphology observed in Rbf-deficient CySCs (Fig. 6H).

Overall, our data show that Rbf-deficient CySCs express lower levels of genes controlling many aspects of oxidative metabolism than control CySCs, and have both reduced numbers and less developed mitochondria. These results suggest that lower metabolic activity in Rbf-deficient cells may contribute to their inability to differentiate and raise the possibility that alterations in metabolism may play a role in normal cyst cell differentiation.

### Endogenous cyst cell differentiation is associated with metabolic gene expression changes

To test whether wild type cyst cell differentiation did indeed involve increased expression of metabolic genes, we compared gene expression profiles in CySCs and differentiating cyst cells in control animals. The CySC pool, presumably also including a fraction of their most recent, immediate progeny, was labelled using the *zfh1-T2A-Gal4* genomic knock-in line (Albert et al., 2018) to express the nuclear dsRed derivative, RedStinger (Fig. S8A). We used the Gal4-Gal80-Hack system (Lin and Potter, 2016) to convert the *zfh1-T2A-Gal4* driver into a *zfh1-T2A-T2A-Gal80* repressor line (see Materials and Methods), which we combined with *tj-Gal4* to label the entire cyst cell lineage with the exception of the Zfh1-positive stem cells (Fig. S8B). Between 125 and 150 dissociated cells per replica were sorted and processed for RNA sequencing. Principal component analysis considering all expressed genes separated the ten transcriptomes into two non-overlapping clusters (Fig. S8C). Applying stringent criteria of FDR<10^−3^ and an absolute of the log_2_(fold change)>1.5 we found 571 upregulated genes in CySCs compared to the cyst cell population and 1284 downregulated ones (Fig. 7A, Supplementary File S1). We examined known CySC and cyst cell markers to validate our results. *zfh1* transcripts were detected at a 5.4-fold higher level in the CySC samples relative to differentiated cyst cells (Fig. S8D and see Supplementary File S1). Consistent with previous observations (Inaba et al., 2011; Li et al., 2003; Schulz et al., 2002; Voog et al., 2014), Zfh1-positive CySCs exhibited 2.2-fold higher expression of *tj*, and 1.6-fold of *string* (*stg)* than Zfh1-negative differentiated cyst cells (Fig. S8D-E, Supplementary File S1). Conversely, expression of the differentiation marker *eya* (Fabrizio et al., 2003) was upregulated 6.2-fold in differentiated cyst cells relative to CySCs, while *Rbf* expression was 12.5-fold higher (Fig. S8D-E).

**Figure 7.**
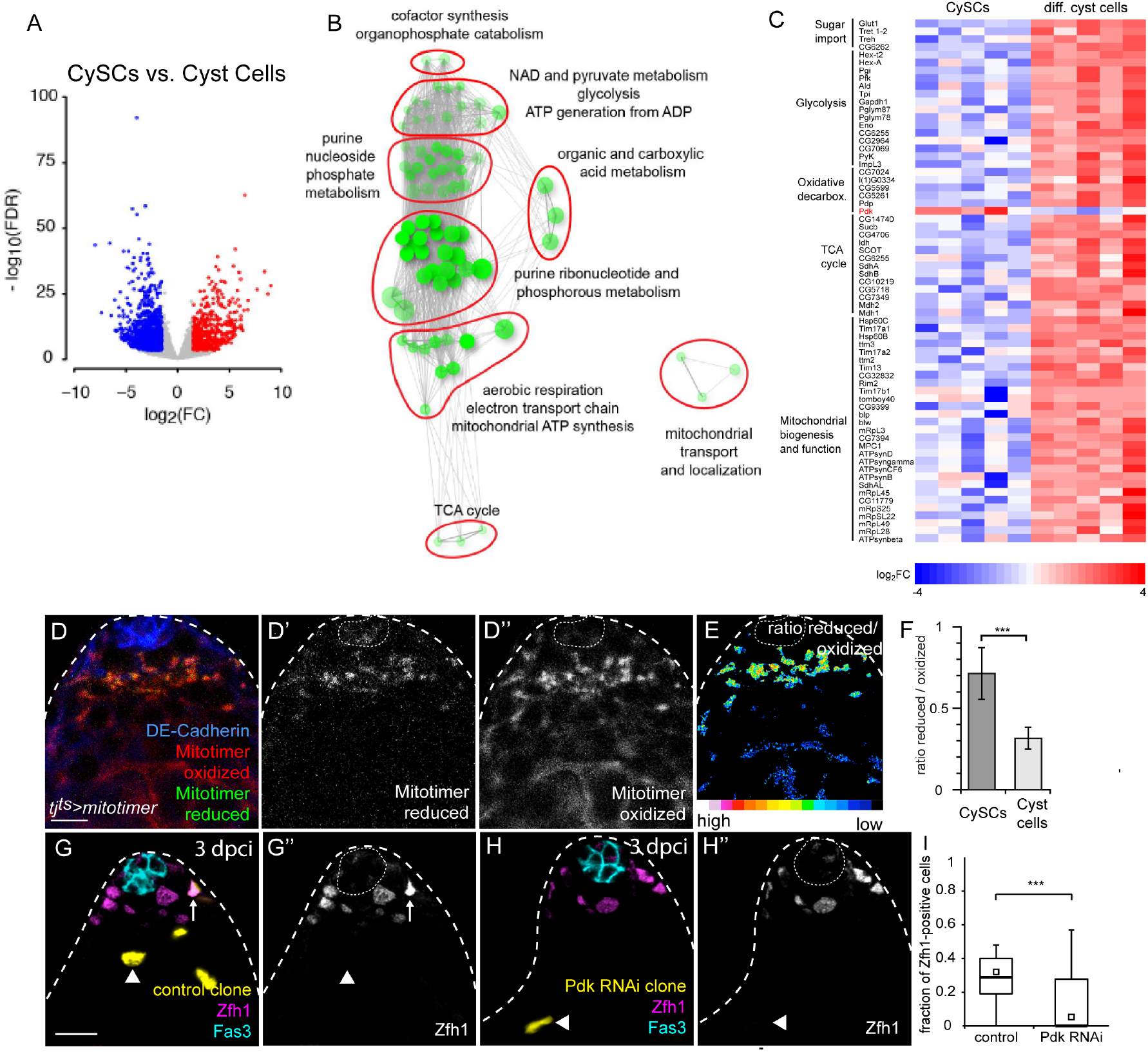
Endogenous cyst cell differentiation involves changes in metabolic gene expression and activity. (A) Volcano plot showing log_10_ of false discovery rate (FDR) against the log_2_(fold change) for individual genes when comparing expression from sorted CySCs and differentiating cyst cells. Genes whose expression was log_2_(fold change)>1.5 downregulated and upregulated with an FDR below 0.001 are labelled in blue and red, respectively. (B) Gene ontology analysis of genes downregulated in CySCs compared to differentiating cyst cells. The size of the circles represents the number of genes in each biological process. A higher significance of enrichment is shown as stronger colouring of the circle. Lines connect nodes with at least 15% of shared genes and their thickness increases with the percentage of shared genes. CySCs express lower levels of genes involved in many aspects of energy generation and mitochondrial function. (C) Heatmap showing relative expression in CySCs versus cyst cells of genes whose products are involved in energy generation from glucose. Colour key shows expression change in log_2_(fold change); blue represents lower expression, while higher expression is shown in red. (D) Expression of the mitochondrial dye Mitotimer, a slow-maturing, mitochondrially-targeted RFP, which transitions from green to red fluorescence through oxidation in the mitochondrial matrix, in a control testis driven with *tj*^*ts*^. Oxidized (red, single channel D”) reporter is detected both in CySCs close to the hub, labelled with DE-Cadherin (blue) and in cyst cells distant from it. However, reduced (green, single channel D’) reporter was mainly present in CySCs surrounding the niche. (E) Ratio of reduced/oxidized mitochondria, showing higher values in CySCs (F) Quantification of the ratio of reduced to oxidized Mitotimer, showing a significantly higher ratio in CySCs compared to mature cyst cells. *** denotes P<0.001, n=19 testes, Student’s t-test. (G,H) Positively-labelled clones at 3 dpci control clones expressing RFP (yellow, single channels in G’,H’), labelled with Zfh1 (magenta, single channels in G”,H”) to identify CySCs and Fas3 (cyan) to label the hub. Control clones (G) were composed of Zfh1-expressing (arrow) and non-expressing cells (arrowhead). Clones expressing Pdk RNAi contained only differentiated cyst cells (arrowhead). Dotted lines outline the hub. Scale bars: 20µm. (I) Quantification of Zfh1 positive cells within each clone at 3 dpci. Clones expressing Pdk RNAi contained a significantly lower fraction of Zfh1-positive CySCs (n=28 for control clones and n=29 for Pdk RNAi clones, Kruskal-Wallis test with Tukey’s HSD post hoc test, *** denotes P<0.001).

Having validated that our approach enabled us to identify genes that were differentially expressed in and relevant to the function of CySCs, we next asked what transcriptional changes occurred during normal cyst cell differentiation. Genes upregulated in cyst cells relative to CySCs were enriched for GO terms associated with cellular metabolism, and could be grouped together into broad categories encompassing ATP generation, NAD and pyruvate metabolism, TCA cycle and mitochondrial transport (Fig. 7B). Further analysis revealed that differentiating cyst cells upregulated genes whose products are responsible for all aspects of energy generation from glucose, including sugar import and glycolysis, TCA cycle components, and mitochondrial biosynthesis and function (Fig. 7C), consistent with the more mature mitochondrial morphology visible in differentiated cyst cells by electron microscopy (Fig. 6E,F). These differences in gene expression suggested that cyst cells and CySCs differed in their cell physiology, in particular that differentiated cyst cells had a metabolic state biased towards increased oxidative phosphorylation. To test this, we used the mitotimer sensor (Laker et al., 2014), which consists of a mitochondrially-targeted RFP that is initially present in an immature green fluorescent precursor but transitions to the mature red fluorescent form when oxidised. The rate of precursor oxidation, as measured by ratiometric imaging of the red and green fluorescence, can be used as a readout for the oxidative state of the mitochondrial matrix in the different cell types. Using *tj*^*ts*^ to express the reporter in the cyst lineage in a single overnight pulse, we observed that the mitochondrial matrix of differentiated cyst cells had a lower ratio of reduced to oxidised mitotimer than CySCs (Fig. 7D-F), indicating increased oxidation in the mitochondrial matrix as cyst cells differentiate.

Next, we asked whether increasing mitochondrial activity in CySCs could induce differentiation. The entry of pyruvate, the product of glycolysis, into mitochondria to feed the TCA cycle is rate-limiting for mitochondrial oxidation. This step is regulated by Pyruvate dehydrogenase, which is antagonised by Pyruvate dehydrogenase kinase (Pdk). *Pdk* expression levels in CySCs were roughly twice as high as in differentiated cyst cells (Fig 7C, Supplementary file S1), while its antagonistic phosphatase Pdp, encoding an activator of mitochondrial Pyruvate consumption, was more highly expressed in cyst cells (Fig 7C, Supplementary file S1), a pattern consistent with lower mitochondrial activity in CySCs than cyst cells. We hypothesised that loss of *Pdk* should result in increased pyruvate availability for the mitochondrial TCA cycle and lead to precocious differentiation. Indeed, knockdown of *Pdk* resulted in rapid loss of CySCs, such that by 3 dpci there were 0.12±0.03 Zfh1-positive cells per clone compared to 0.31±0.04 in controls (Kruskal-Wallis test with Tukey’s HSD post hoc test, P<0.001, n=28 and n=27, respectively) (Fig. 7G-I). Thus, knocking down a negative regulator of mitochondrial pyruvate consumption is sufficient to drive differentiation in the cyst lineage.

Altogether, we show that differentiation of CySCs into cyst cells involves significant transcriptional changes of genes regulating cellular metabolism, resulting in differences in mitochondrial activity during differentiation and that these shifts in metabolic activity are sufficient to drive changes in cell identity.

### Promoting mitochondrial biogenesis can rescue differentiation but not cell cycle exit in Rbf knockdown testes

Our results indicate that oxidative metabolism increases during cyst cell differentiation, and that, conversely, CySCs lacking Rbf have decreased expression of genes encoding mitochondrial proteins and fewer mitochondria. We asked how similar the gene expression profile of Rbf-deficient CySCs was to wild-type CySCs. Approximately 27% of genes that were differentially expressed between CySCs and cyst cells were similarly changed in Rbf-deficient CySCs (Fig. S9A, Supplementary file S1). Additionally, GO analysis of the overlapping downregulated genes revealed enrichment for categories related to oxidation-reduction and carboxylic acid metabolic processes (Fig. S9B). Genes downregulated in these categories include glycolytic and mitochondrial enzymes, such as *Ldh, PyK, Gapdh, blw* or *Mtp*α. These data suggest that the gene expression of Rbf-deficient CySCs is similar to wild type CySCs for metabolic genes and that the metabolic state of Rbf-deficient CySCs may be limiting their ability to differentiate.

Since Rbf-deficient CySCs showed decreased expression of metabolic genes and fewer mitochondria, and since increased mitochondrial activity is implicated in cyst cell differentiation, we asked whether promoting oxidative respiration in Rbf-deficient CySCs could rescue their block in differentiation. To increase mitochondrial biogenesis and activity in Rbf-deficient cells, we expressed the transcription factor Spargel (Srl, homologue of PGC1α), which, together with its partner Delg (Flybase: Ets97D, the homologue of NRF-2α), regulates expression of mitochondrial genes and promotes mitochondrial biogenesis and activity (Rera et al., 2011; Tiefenbock et al., 2010).

While knockdown of Rbf resulted in expansion of Zfh1 expression away from the hub and a lack of Eya-expressing cells (Figs. 8A,B), co-expression of Srl and Delg led to a distribution of cell types that resembled control testes: Zfh1-expressing cells were restricted to about two rows surrounding the hub, while cells further away expressed Eya (Fig. 8C). Similarly, clones expressing Rbf RNAi together with either Srl or Delg were partially rescued in their ability to differentiate (Fig. S10), and contained Eya-expressing cyst cells in 47% (n=36) and 26% (n=35) of cases, respectively, compared to 9% in Rbf knockdown alone (n=46). Although the increase of differentiated cells in Delg-overexpressing clones was not significant (Fisher’s exact test, P=0.06), overexpression of Srl resulted in a statistically significant rescue of differentiation (Fisher’s exact test, P<0.0001). Compared to controls, the rescued Eya-positive cells appeared halted in their differentiation: they did not express as high levels of Eya and their nuclei did not grow as large as those of wild type differentiated cyst cells (Fig. 8A and 8C). When we assayed for proliferation in the rescued testes, we observed Eya-positive, Zfh1-negative cells that were also EdU-positive (Fig. 8C, arrowheads) in 5/7 of testes, which we never observed in controls (0/16).

**Figure 8.**
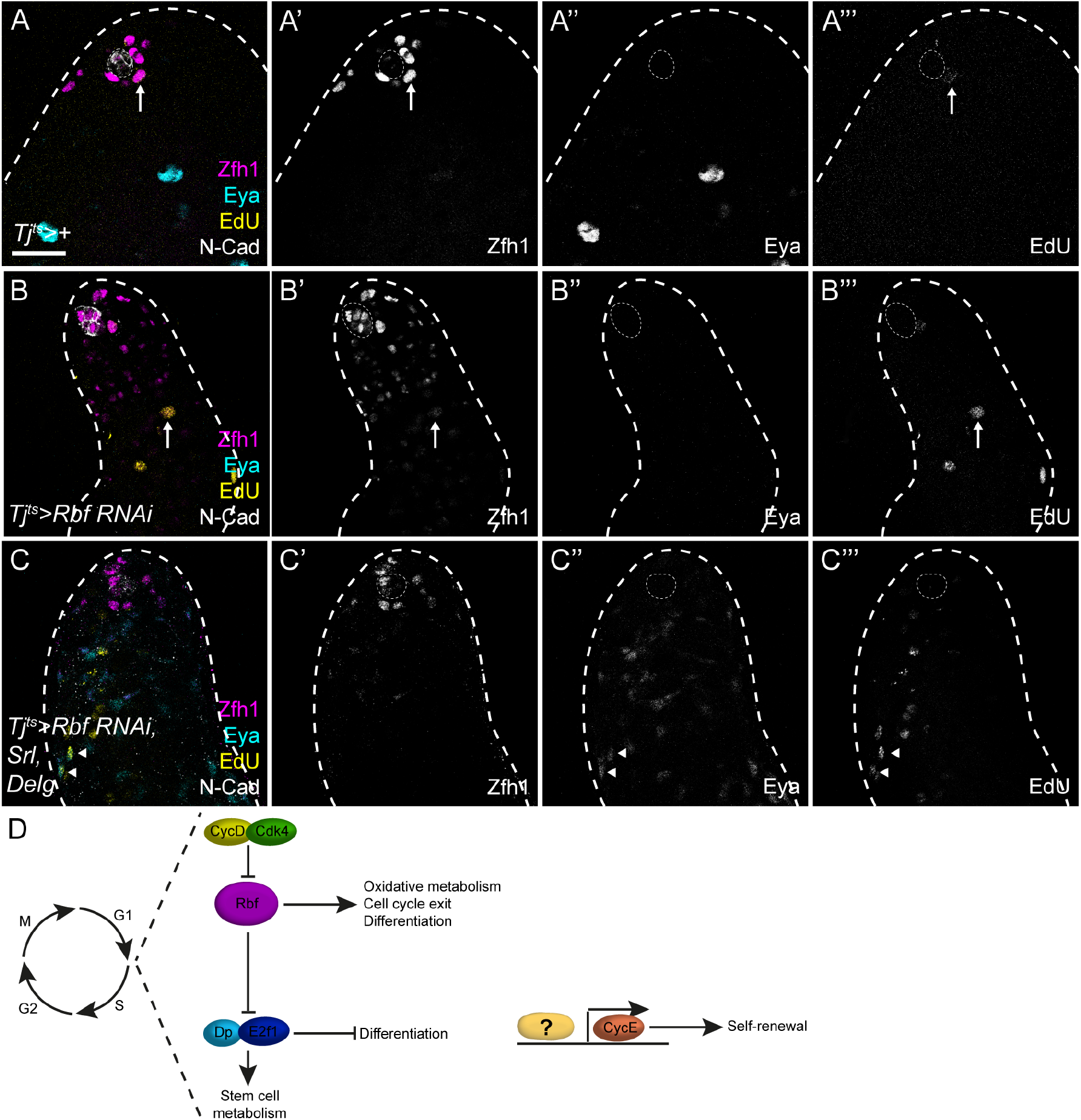
Promoting mitochondrial biogenesis can rescue differentiation but not cell cycle exit in Rbf knockdown testes. (A-C) Testes labelled with Zfh1 (magenta, single channel in A’,B’,C’), Eya (cyan, single channel in A”,B”,C”) EdU (yellow, single channel in A’”,B’”,C’”), and N-Cadherin (N-Cad) to identify the hub. (A) In controls, Zfh1 expression is detected only in cells surrounding the hub and EdU incorporation in somatic cells is restricted to Zfh1-expressing cells. (B) Knockdown of Rbf results in ectopic Zfh1 expression and a complete absence of Eya expression. EdU-positive proliferating cells can be seen distant from the hub (arrow). (C) Mis-expression of Srl and Delg together with Rbf knockdown restores Eya expression in somatic cells away from the hub, while Zfh1 is restricted to cells surrounding the hub. EdU-positive, Eya-positive cells (arrowheads) are visible away from the hub, indicating that differentiated cells no longer exit the cell cycle. Dotted lines outline the hub. Scale bars: 20µm. (D) Diagram summarising the regulation of cell cycle progression and self-renewal in CySCs.

Thus, increasing mitochondrial activity rescues the ability of Rbf-deficient CysCs to differentiate but not to exit the cell cycle. These data suggest that coordinating cell cycle exit and differentiation in cyst cells is achieved by Rbf-dependent inhibition of E2f1/Dp activity, while ectopic E2f1/Dp promotes a metabolic state that inhibits differentiation.

## Discussion

In this study, we asked whether cell cycle regulators affected the decision of stem cells to self-renew or differentiate in the Drosophila testis. We show that continued cycling and maintenance of CySC identity depend on expression of the G1/S cyclin *cycE*, but the E2f1/Dp transcription factor which in other contexts is thought to regulate *cycE* expression, is dispensable for CySC self-renewal, indicating an unusual regulation of S-phase entry in CySCs (Fig. 8D). In contrast, loss of the negative regulator of G1/S transition, Rbf, prevents both cell cycle exit and differentiation, acting entirely through regulation of E2f1/Dp transcriptional activity. Differentiation therefore endogenously necessitates a silencing of E2f1/Dp. We further show that ectopic E2f1/Dp activity results in reduced expression of genes encoding metabolic enzymes, and that normal cyst cell differentiation involves upregulation of metabolic gene expression and of mitochondrial oxidation. Finally, promoting mitochondrial biogenesis is sufficient to rescue the differentiation of Rbf-deficient cyst cells, but not their exit from the cell cycle. Thus, we conclude that Rbf coordinates differentiation and cell cycle exit in G1 by silencing E2f1/Dp activity and enabling a metabolic state compatible with differentiation (Fig. 8D).

Our observations indicate that the regulatory networks controlling S phase entry in CySCs differ from those described in other cells. While *cycE* is necessary for CySC proliferation, *E2f1* and *Dp* are dispensable. *cycE* expression is thought to be transcriptionally induced by E2f1/Dp and we note that ectopic E2f1/Dp is able to induce *cycE* expression, as *cycE* transcripts were upregulated in Rbf-deficient testes (Fig. 5C), and ectopic CycE was observed in larval testes mutant for *Rbf* (Dominado et al., 2016). However, continued proliferation in *Dp* or *E2f1* mutant CySCs indicates that this regulation is not necessary and that other inputs may impinge on *cycE* regulation. One possibility is that self-renewal signals induce *cycE* expression, and indeed, two signals known to be active in CySCs and required for their self-renewal, Hedgehog and Hippo, are known regulators of *cycE* (Amoyel et al., 2013; Amoyel et al., 2014; Duman-Scheel et al., 2002; Huang et al., 2005; Michel et al., 2012). In particular, Zfh1 directly inhibits Hippo activity in CySCs, restricting Yki activation to the CySC pool, and suggesting a possible link to *cycE* expression (Albert et al., 2018). It is intriguing to note that in female GSCs, CycE is detected in G2 and M phases as well as G1, indicating that its expression may be regulated differently in different cell types (Hsu et al., 2008).

We find that despite a reporter pattern consistent with periodic cell cycle-dependent activation, E2f1/Dp are not required for normal cycling in CySCs. While surprising, this result is consistent with findings that *Dp* null and *E2f1*/*E2f2* double mutants are viable until pupal stages (Frolov et al., 2001), suggesting that most larval proliferation can occur without Dp activity. Indeed, recent work showed that *Dp* null mutants could be rescued to adulthood if Dp was restored only in muscle using the Mef2-Gal4 driver (Zappia and Frolov, 2016). Similarly, studies in the mouse retina showed that retinal progenitors and Müller glia deficient for all three mammalian activator E2fs could continue to proliferate (Chen et al., 2009). Our data instead argue that the role of E2f1/Dp activity is to promote a metabolic state that prevents differentiation. Thus, restraining E2f1/Dp activity through Rbf is essential to allow cyst cell differentiation, such that both differentiation and cell cycle exit are coordinated through regulation of E2f/Dp. In the best-characterised examples, Rb is typically thought to influence differentiation through interactions with tissue-specific transcription factors (Chen et al., 1996; Gu et al., 1993; Sellers et al., 1998; Thomas et al., 2001); however in the mouse retina, cell cycle exit and differentiation of amacrine cells separately depends on Rb-mediated inhibition of two activator E2fs, E2f1, and E2f3a, respectively (Chen et al., 2007). Our results indicate that E2f’s functions in antagonising differentiation may be conserved. Indeed, Rb and E2f/Dp regulate metabolism in mice and Drosophila (Blanchet et al., 2011; Guarner et al., 2017; Nicolay et al., 2015; Zappia et al., 2019). Intriguingly, Rbf, E2f1, E2f2 and Dp directly bind the enhancers of several genes encoding mitochondrial-associated proteins both in Drosophila larvae and mammalian cells (Ambrus et al., 2013; Blanchet et al., 2011). However, the mechanisms of action appear cell-specific as in larval Drosophila tissues, E2f and Dp maintain mitochondrial gene expression and activity, while in differentiated skeletal muscle, E2f1 acts together with Rb to inhibit oxidative metabolic gene expression. This contrasts with the situation in the CySCs, where many genes encoding mitochondrial factors have reduced expression upon ectopic E2f1/Dp activity, in a manner antagonised by Rbf. These observations suggest that in the testis the regulation may be indirect. In many instances, Rb loss results in dysregulation of chromatin regulators, suggesting a potential mechanism by which these effects could be mediated (Benevolenskaya et al., 2005; Gonzalo et al., 2005).

Overall, our results suggest a model for how cell cycle exit and differentiation are linked in CySCs: by limiting the ability of cells to change their metabolic state, E2f1/Dp activity ensures that cycling cells cannot differentiate.

## Methods

### Fly stocks and husbandry

Lineage-wide misexpression and knockdown experiments were carried out using the *tj-Gal4* driver, together with a *Tub>Gal80*^*ts*^ transgene (referred to as *tj*^*ts*^) to control the temporal pattern of expression (McGuire et al., 2004). Crosses were raised at 18°C. Males were collected 0-3 days after eclosion and shifted to 29°C for 10 days. The following stocks were used: *UAS-Rbf RNAi* (BDSC #41863); *UAS-CycE* (BDSC #4781); *UAS-Dp RNAi* (VDRC #12722); *PCNA-GFP; UAS-E2f1, UAS-Dp* (gifts of L. Buttitta); *Ald1-GFP* (Kyoto DGRC #115279); *UAS-mitotimer* (BDSC #57323); *UAS-Srl* (gift of H. Stocker); *UAS-Delg* (gift of M. Simonelig); *esg-GFP* (gift of L. Jones).

*w;; zfh1-T2A-T2A-Gal80* (referred to in brief as *zfh1-Gal80*) was generated by crossing yw vasa-Cas9 first to zfh1-T2A-Gal4 w+/TM3, Sb (Albert et al., 2018) and then to the Pin/CyO; Gal4-Gal80Hack/TM6B (94E5) Gal4-Gal80 HACK stock (Lin and Potter, 2016) that contains all the required components such as gRNA genes, homology arms, and an eye RFP selection marker to insert a T2A-Gal80 cassette into the Gal4 ORF of any Gal4 transgene on the homologous chromosome.

For clonal analysis, flies were raised and maintained at 25°C. Adult flies were collected 0-3 days after eclosion and heat shocked at 37°C for 1 hour. Negatively marked Rbf mutant clones were generated using a duplication of the X chromosome on the third chromosome, *Dp(1:3)DC012* (Venken et al., 2010), which fully rescues the viability of *Rbf*^*14*^ hemizygous mutants. The experimental genotype was *Rbf*^*14*^ *w/Y; hs-flp/+; ubi-GFP Dp(1:3)DC012 FRT*^*2A*^*/FRT*^*2A*^. Control clones were generated in the same way but wild type for *Rbf* in the endogenous locus (*y,w,hsflp*^*122*^*/Y;; ubi-GFP Dp(1:3)DC012 FRT*^*2A*^*/FRT*^*2A*^).

All other clones were generated by the MARCM technique (Lee and Luo, 2001). Stocks used to generate clones were: *y,w,hsflp*^*122*,^*Tub>Gal4,UAS-nlsGFP; FRT*^*42D*^,*Tub>Gal80,CD71*; *y,w,hsflp*^*122*^,*Tub>Gal4,UAS-nlsGFP;FRT*^*40A*^,*Tub>Gal80; y,w,hsflp*^*122*^,*Tub>Gal4,UAS-nlsGFP;; FRT*^*82B*^,*Tub>Gal80* and *w,hs-FLP,C587-Gal4,UAS-RedStinger; FRT*^*42D*^,*Tub>Gal80*. We used the following alleles: *CycE*^*AR95*^ (gift of A. Audibert); *E2f1*^*rM729*^ (also known as *E2f1*^*729*^, BDSC#35849); *Dpa3* and *Dp*^*a4*^ (gift of M. Frolov). *Dp* and *Ef21* mutants were validated by lack of complementation against *Df(2R)Exel7124* (BDSC #7872), and *Df(3R)Exel6186* (BDSC #7665), respectively, and in addition, the *Dp*^*a3*^ hemizygous mutant was rescued to adulthood by Dp over-expression with *Mef2-Gal4*, as previously described (Zappia and Frolov, 2016). We also used the constructs *UAS-Rbf RNAi* (BDSC #41863); *UAS-Rbf RNAi* (BDSC #36744); *UAS-Pdk RNAi* (BDSC #28635); *UAS-Srl* (gift of H. Stocker); *UAS-Delg* (gift of M. Simonelig).

### Immunohistochemistry

The following antibodies were used: rat anti-Chinmo (gift of N.Sokol), 1:50; mouse anti-Eya (Developmental Studies Hybridoma Bank, DSHB), 1:20; mouse anti-Fas3 (DSHB), 1:20; mouse anti-Wg (DSHB), 1:500; chicken anti-GFP (Aves Lab, GFP-1010), 1:500; rabbit anti-GFP (Thermo Fisher, A6455), 1:500; rat anti-NCad (DSHB), 1:20; mouse anti-Rbf (gift of N. Dyson), 1:15; guinea pig anti-Tj (gift of D. Godt), 1:3000; rabbit anti-Zfh1 (gift of R. Lehmann), 1:5000. Fixing and immunohistochemistry was carried out as previously described (Flaherty et al., 2010; Michel et al., 2011).

For EdU staining, testes were dissected in Schneider’s medium and incubated for 30 minutes at room temperature in Schneider’s medium containing 10µM EdU. Samples were then fixed and incubated with primary and secondary antibodies as above. Click reaction was then carried out for 30 minutes at room temperature in the following reaction buffer: 2.5µM Alexa-405 picolyl azide (Click Chemistry Tools), 0.1mM THPTA, 2mM sodium ascorbate and 1mM CuSO_4_.

### Statistics

Statistical tests were carried out using GraphPad Prism. For comparison of clone recovery rates, Fisher’s exact test was used, for all other experiments, the test is indicated. Numbers shown are mean ± SEM.

### RNAseq experiments

CySCs and differentiating cyst cells were labelled with RedStinger expression driven by *zfh1-T2A-Gal4* and *tj-gal4; zfh1-Gal80*, respectively. In the case of Rbf knockdown experiment, somatic cells were labelled with GFP driven by *tj-gal4*. Testes were dissected in Schneider’s medium and then separation buffer containing Schneider’s medium, collagenase, trypsin and EDTA was added. Samples were vigorously agitated for 15-30 min. The cell suspension was then filtered using a cell strainer.

mRNA was isolated from the samples, reverse transcribed, and amplified using the SmartSeq2 kit (Illumina) for the study within the somatic lineage and SMART-Seq v4 Ultra (Takara Bio) for the Rbf knockdown experiment. Libraries were then generated using the Nextera kit (Illumina) and 75bp single end read-sequencing was carried out.

Reads were quality checked and mapped using the A.I.R. RNAseq web-based analysis package (Sequentia Biotech, Barcelona). The underlying algorithms and software packages collated in A.I.R. are described in (Vara et al., 2019). To assign differential gene expression between samples we used the DESeq algorithm as implemented in the A.I.R. package. In the case of the Rbf knockdown experiment, reads were mapped and aligned using Hisat2 and StringTie. Differential expression was analysed using DESeq2. Volcano plots and heatmaps were generated using DEBrowser (Kucukural et al., 2019) in RStudio. Gene ontology analysis was performed using the shinyGO web platform for both experiments (Ge et al., 2020).

The corresponding raw datasets for the RNAseq experiments have been deposited online and will be available upon publication. Differentially-expressed genes are listed in the supplementary information (Supplementary file S1).

### Electron microscopy

*tj*^*ts*^ crosses were kept at 18°C. Flies 0-4 days after eclosion were collected and shifted to 29°C. Testes were dissected and fixed in 1% glutaraldehyde/paraformaldehyde for 1.5 hours. Samples were then embedded in paraffin and sectioned for imaging.

## Acknowledgements

We would like to thank members of the fly community for fly stocks and reagents, Andrew Herman and Lorena Sueiro Ballesteros at the University of Bristol and Gavin Giel and the Flow Cytometry Core Facility at Philipps-University Marburg for FACS, Christy Waterfall and the Bristol Genomics Facility and Andreas Dahl and Susanne Reinhardt at the CRTD Deep Sequencing Core Facility for the RNA sequencing. Sabina Huhn and Ljubinka Cigoja provided essential technical assistance in fly genetics, molecular cloning and immunohistochemistry. Many thanks to members of the Fernandes, Poole and Barrios labs for constructive discussions, and to Alex Gould, Parthive Patel, Laura Johnston, and Vilaiwan Fernandes for critical feedback on the manuscript.

This work was supported by an MRC Career Development Award MR/P009646/2 to MA and a DFG grant BO 3270/3-1 to CB.

## Author contributions

DSdlM, SH-F, LP, CB and MA conducted experiments and analysed the data. Experiments were conceptualised and designed by DSdlM, MA and CB. DSdlM and MA wrote the manuscript, with editing contributions from CB. All authors contributed to the article and approved the submitted version.

## Notes

### Competing Interest Statement

The authors have declared no competing interest.

## References

Ables, E. T. and Drummond-Barbosa, D. (2013). Cyclin E controls Drosophila female germline stem cell maintenance independently of its role in proliferation by modulating responsiveness to niche signals. Development 140, 530–540.

Albert, E. A., Puretskaia, O. A., Terekhanova, N. V., Labudina, A. and Bokel, C. (2018). Direct control of somatic stem cell proliferation factors by the Drosophila testis stem cell niche. Development 145.

Ambrus, A. M., Islam, A. B., Holmes, K. B., Moon, N. S., Lopez-Bigas, N., Benevolenskaya, E. V. and Frolov, M. V. (2013). Loss of dE2F compromises mitochondrial function. Dev Cell 27, 438–451.

Amoyel, M., Anderson, J., Suisse, A., Glasner, J. and Bach, E. A. (2016). Socs36E Controls Niche Competition by Repressing MAPK Signaling in the Drosophila Testis. PLoS Genet 12, e1005815.

Amoyel, M., Sanny, J., Burel, M. and Bach, E. A. (2013). Hedgehog is required for CySC self-renewal but does not contribute to the GSC niche in the Drosophila testis. Development 140, 56–65.

Amoyel, M., Simons, B. D. and Bach, E. A. (2014). Neutral competition of stem cells is skewed by proliferative changes downstream of Hh and Hpo. EMBO J 33, 2295–2313.

Benevolenskaya, E. V., Murray, H. L., Branton, P., Young, R. A. and Kaelin, W. G., Jr. (2005). Binding of pRB to the PHD protein RBP2 promotes cellular differentiation. Mol Cell 18, 623–635.

Blanchet, E., Annicotte, J. S., Lagarrigue, S., Aguilar, V., Clape, C., Chavey, C., Fritz, V., Casas, F., Apparailly, F., Auwerx, J., et al. (2011). E2F transcription factor-1 regulates oxidative metabolism. Nat Cell Biol 13, 1146–1152.

Brehm, A., Miska, E. A., McCance, D. J., Reid, J. L., Bannister, A. J. and Kouzarides, T. (1998). Retinoblastoma protein recruits histone deacetylase to repress transcription. Nature 391, 597–601.

Buttitta, L. A. and Edgar, B. A. (2007). Mechanisms controlling cell cycle exit upon terminal differentiation. Curr Opin Cell Biol 19, 697–704.

Chen, D., Opavsky, R., Pacal, M., Tanimoto, N., Wenzel, P., Seeliger, M. W., Leone, G. and Bremner, R. (2007). Rb-mediated neuronal differentiation through cell-cycle-independent regulation of E2f3a. PLoS Biol 5, e179.

Chen, D., Pacal, M., Wenzel, P., Knoepfler, P. S., Leone, G. and Bremner, R. (2009). Division and apoptosis of E2f-deficient retinal progenitors. Nature 462, 925–929.

Chen, P. L., Riley, D. J., Chen, Y. and Lee, W. H. (1996). Retinoblastoma protein positively regulates terminal adipocyte differentiation through direct interaction with C/EBPs. Genes Dev 10, 2794–2804.

Cheng, J., Tiyaboonchai, A., Yamashita, Y. M. and Hunt, A. J. (2011). Asymmetric division of cyst stem cells in Drosophila testis is ensured by anaphase spindle repositioning. Development 138, 831–837.

Cheng, T., Rodrigues, N., Shen, H., Yang, Y., Dombkowski, D., Sykes, M. and Scadden, D. T. (2000). Hematopoietic stem cell quiescence maintained by p21cip1/waf1. Science 287, 1804–1808.

DeGregori, J., Kowalik, T. and Nevins, J. R. (1995). Cellular targets for activation by the E2F1 transcription factor include DNA synthesis- and G1/S-regulatory genes. Mol Cell Biol 15, 4215–4224.

Dimova, D. K. and Dyson, N. J. (2005). The E2F transcriptional network: old acquaintances with new faces. Oncogene 24, 2810–2826.

Dimova, D. K., Stevaux, O., Frolov, M. V. and Dyson, N. J. (2003). Cell cycle-dependent and cell cycle-independent control of transcription by the Drosophila E2F/RB pathway. Genes Dev. 17, 2308–2320.

Dominado, N., La Marca, J. E., Siddall, N. A., Heaney, J., Tran, M., Cai, Y., Yu, F., Wang, H., Somers, W. G., Quinn, L. M., et al. (2016). Rbf Regulates Drosophila Spermatogenesis via Control of Somatic Stem and Progenitor Cell Fate in the Larval Testis. Stem Cell Reports 7, 1152–1163.

Dubey, P., Kapoor, T., Gupta, S., Shirolikar, S. and Ray, K. (2019). Atypical septate junctions maintain the somatic enclosure around maturing spermatids and prevent premature sperm release in Drosophila testis. Biol Open 8.

Duman-Scheel, M., Weng, L., Xin, S. and Du, W. (2002). Hedgehog regulates cell growth and proliferation by inducing Cyclin D and Cyclin E. Nature 417, 299–304.

Duronio, R. J., Brook, A., Dyson, N. and O’Farrell, P. H. (1996). E2F-induced S phase requires cyclin E. Genes Dev. 10, 2505–2513.

Duronio, R. J., O’Farrell, P. H., Xie, J. E., Brook, A. and Dyson, N. (1995). The transcription factor E2F is required for S phase during Drosophila embryogenesis. Genes Dev 9, 1445–1455.

Dynlacht, B. D., Brook, A., Dembski, M., Yenush, L. and Dyson, N. (1994). DNA-binding and trans-activation properties of Drosophila E2F and DP proteins. Proc Natl Acad Sci U S A 91, 6359–6363.

Fabrizio, J. J., Boyle, M. and DiNardo, S. (2003). A somatic role for eyes absent (eya) and sine oculis (so) in Drosophila spermatocyte development. Dev Biol 258, 117–128.

Fairchild, M. J., Islam, F. and Tanentzapf, G. (2017). Identification of genetic networks that act in the somatic cells of the testis to mediate the developmental program of spermatogenesis. PLoS Genet 13, e1007026.

Fairchild, M. J., Smendziuk, C. M. and Tanentzapf, G. (2015). A somatic permeability barrier around the germline is essential for Drosophila spermatogenesis. Development 142, 268–281.

Flaherty, M. S., Salis, P., Evans, C. J., Ekas, L. A., Marouf, A., Zavadil, J., Banerjee, U. and Bach, E. A. (2010). chinmo is a functional effector of the JAK/STAT pathway that regulates eye development, tumor formation, and stem cell self-renewal in Drosophila. Dev Cell 18, 556–568.

Frolov, M. V., Huen, D. S., Stevaux, O., Dimova, D., Balczarek-Strang, K., Elsdon, M. and Dyson, N. J. (2001). Functional antagonism between E2F family members. Genes Dev 15, 2146–2160.

Frolov, M. V., Moon, N. S. and Dyson, N. J. (2005). dDP is needed for normal cell proliferation. Mol Cell Biol 25, 3027–3039.

Ge, S. X., Jung, D. and Yao, R. (2020). ShinyGO: a graphical gene-set enrichment tool for animals and plants. Bioinformatics 36, 2628–2629.

Gonczy, P. and DiNardo, S. (1996). The germ line regulates somatic cyst cell proliferation and fate during Drosophila spermatogenesis. Development 122, 2437–2447.

Gonzalo, S., Garcia-Cao, M., Fraga, M. F., Schotta, G., Peters, A. H., Cotter, S. E., Eguia, R., Dean, D. C., Esteller, M., Jenuwein, T., et al. (2005). Role of the RB1 family in stabilizing histone methylation at constitutive heterochromatin. Nat Cell Biol 7, 420–428.

Greenspan, L. J. and Matunis, E. L. (2018). Retinoblastoma Intrinsically Regulates Niche Cell Quiescence, Identity, and Niche Number in the Adult Drosophila Testis. Cell Rep 24, 3466–3476 e3468.

Gu, W., Schneider, J. W., Condorelli, G., Kaushal, S., Mahdavi, V. and Nadal-Ginard, B. (1993). Interaction of myogenic factors and the retinoblastoma protein mediates muscle cell commitment and differentiation. Cell 72, 309–324.

Guarner, A., Morris, R., Korenjak, M., Boukhali, M., Zappia, M. P., Van Rechem, C., Whetstine, J. R., Ramaswamy, S., Zou, L., Frolov, M. V., et al. (2017). E2F/DP Prevents Cell-Cycle Progression in Endocycling Fat Body Cells by Suppressing dATM Expression. Dev Cell 43, 689–703 e685.

Hardy, R. W., Tokuyasu, K. T., Lindsley, D. L. and Garavito, M. (1979). The germinal proliferation center in the testis of Drosophila melanogaster. J Ultrastruct Res 69, 180–190.

Hsu, H. J., LaFever, L. and Drummond-Barbosa, D. (2008). Diet controls normal and tumorous germline stem cells via insulin-dependent and -independent mechanisms in Drosophila. Dev Biol 313, 700–712.

Huang, J., Wu, S., Barrera, J., Matthews, K. and Pan, D. (2005). The Hippo signaling pathway coordinately regulates cell proliferation and apoptosis by inactivating Yorkie, the Drosophila Homolog of YAP. Cell 122, 421–434.

Inaba, M., Yuan, H. and Yamashita, Y. M. (2011). String (Cdc25) regulates stem cell maintenance, proliferation and aging in Drosophila testis. Development 138, 5079–5086.

Ishida, S., Huang, E., Zuzan, H., Spang, R., Leone, G., West, M. and Nevins, J. R. (2001). Role for E2F in control of both DNA replication and mitotic functions as revealed from DNA microarray analysis. Mol Cell Biol 21, 4684–4699.

Jones, D. L. and Wagers, A. J. (2008). No place like home: anatomy and function of the stem cell niche. Nat Rev Mol Cell Biol 9, 11–21.

Kato, J., Matsushime, H., Hiebert, S. W., Ewen, M. E. and Sherr, C. J. (1993). Direct binding of cyclin D to the retinoblastoma gene product (pRb) and pRb phosphorylation by the cyclin D-dependent kinase CDK4. Genes Dev 7, 331–342.

Kiger, A. A., White-Cooper, H. and Fuller, M. T. (2000). Somatic support cells restrict germline stem cell self-renewal and promote differentiation. Nature 407, 750–754.

Korenjak, M., Anderssen, E., Ramaswamy, S., Whetstine, J. R. and Dyson, N. J. (2012). RBF binding to both canonical E2F targets and noncanonical targets depends on functional dE2F/dDP complexes. Mol Cell Biol 32, 4375–4387.

Kucukural, A., Yukselen, O., Ozata, D. M., Moore, M. J. and Garber, M. (2019). DEBrowser: interactive differential expression analysis and visualization tool for count data. BMC Genomics 20, 6.

Laker, R. C., Xu, P., Ryall, K. A., Sujkowski, A., Kenwood, B. M., Chain, K. H., Zhang, M., Royal, M. A., Hoehn, K. L., Driscoll, M., et al. (2014). A novel MitoTimer reporter gene for mitochondrial content, structure, stress, and damage in vivo. J Biol Chem 289, 12005–12015.

Leatherman, J. L. and Dinardo, S. (2008). Zfh-1 controls somatic stem cell self-renewal in the Drosophila testis and nonautonomously influences germline stem cell self-renewal. Cell Stem Cell 3, 44–54.

Lee, T. and Luo, L. (2001). Mosaic analysis with a repressible cell marker (MARCM) for Drosophila neural development. Trends Neurosci 24, 251–254.

Li, M. A., Alls, J. D., Avancini, R. M., Koo, K. and Godt, D. (2003). The large Maf factor Traffic Jam controls gonad morphogenesis in Drosophila. Nat Cell Biol 5, 994–1000.

Lin, C. C. and Potter, C. J. (2016). Editing Transgenic DNA Components by Inducible Gene Replacement in Drosophila melanogaster. Genetics 203, 1613–1628.

Matsushime, H., Quelle, D. E., Shurtleff, S. A., Shibuya, M., Sherr, C. J. and Kato, J. Y. (1994). D-type cyclin-dependent kinase activity in mammalian cells. Mol Cell Biol 14, 2066–2076.

McGuire, S. E., Mao, Z. and Davis, R. L. (2004). Spatiotemporal gene expression targeting with the TARGET and gene-switch systems in Drosophila. Sci STKE 2004, pl6.

Michel, M., Kupinski, A. P., Raabe, I. and Bokel, C. (2012). Hh signalling is essential for somatic stem cell maintenance in the Drosophila testis niche. Development 139, 2663–2669.

Michel, M., Raabe, I., Kupinski, A. P., Perez-Palencia, R. and Bokel, C. (2011). Local BMP receptor activation at adherens junctions in the Drosophila germline stem cell niche. Nat Commun 2, 415.

Nicolay, B. N., Danielian, P. S., Kottakis, F., Lapek, J. D., Jr., Sanidas, I., Miles, W. O., Dehnad, M., Tschop, K., Gierut, J. J., Manning, A. L., et al. (2015). Proteomic analysis of pRb loss highlights a signature of decreased mitochondrial oxidative phosphorylation. Genes Dev 29, 1875–1889.

Ohtsubo, M., Theodoras, A. M., Schumacher, J., Roberts, J. M. and Pagano, M. (1995). Human cyclin E, a nuclear protein essential for the G1-to-S phase transition. Mol Cell Biol 15, 2612–2624.

Papagiannouli, F., Berry, C. W. and Fuller, M. T. (2019). The Dlg Module and Clathrin-Mediated Endocytosis Regulate EGFR Signaling and Cyst Cell-Germline Coordination in the Drosophila Testis. Stem Cell Reports 12, 1024–1040.

Pardee, A. B. (1974). A restriction point for control of normal animal cell proliferation. Proc Natl Acad Sci U S A 71, 1286–1290.

Rera, M., Bahadorani, S., Cho, J., Koehler, C. L., Ulgherait, M., Hur, J. H., Ansari, W. S., Lo, T., Jr., Jones, D. L. and Walker, D. W. (2011). Modulation of longevity and tissue homeostasis by the Drosophila PGC-1 homolog. Cell Metab 14, 623–634.

Royzman, I., Whittaker, A. J. and Orr-Weaver, T. L. (1997). Mutations in Drosophila DP and E2F distinguish G1-S progression from an associated transcriptional program. Genes Dev 11, 1999–2011.

Ruijtenberg, S. and van den Heuvel, S. (2016). Coordinating cell proliferation and differentiation: Antagonism between cell cycle regulators and cell type-specific gene expression. Cell Cycle 15, 196–212.

Sage, J. (2012). The retinoblastoma tumor suppressor and stem cell biology. Genes Dev 26, 1409–1420.

Sawado, T., Yamaguchi, M., Nishimoto, Y., Ohno, K., Sakaguchi, K. and Matsukage, A. (1998). dE2F2, a novel E2F-family transcription factor in Drosophila melanogaster. Biochem Biophys Res Commun 251, 409–415.

Schulz, C., Wood, C. G., Jones, D. L., Tazuke, S. I. and Fuller, M. T. (2002). Signaling from germ cells mediated by the rhomboid homolog stet organizes encapsulation by somatic support cells. Development 129, 4523–4534.

Sellers, W. R., Novitch, B. G., Miyake, S., Heith, A., Otterson, G. A., Kaye, F. J., Lassar, A. B. and Kaelin, W. G., Jr. (1998). Stable binding to E2F is not required for the retinoblastoma protein to activate transcription, promote differentiation, and suppress tumor cell growth. Genes Dev 12, 95–106.

Stevaux, O. and Dyson, N. J. (2002). A revised picture of the E2F transcriptional network and RB function. Curr. Opin. Cell Biol. 14, 684–691.

Thacker, S. A., Bonnette, P. C. and Duronio, R. J. (2003). The Contribution of E2F-Regulated Transcription to Drosophila PCNA Gene Function. Curr Biol 13, 53–58.

Thomas, D. M., Carty, S. A., Piscopo, D. M., Lee, J. S., Wang, W. F., Forrester, W. C. and Hinds, P. W. (2001). The retinoblastoma protein acts as a transcriptional coactivator required for osteogenic differentiation. Mol Cell 8, 303–316.

Tiefenbock, S. K., Baltzer, C., Egli, N. A. and Frei, C. (2010). The Drosophila PGC-1 homologue Spargel coordinates mitochondrial activity to insulin signalling. EMBO J 29, 171–183.

Tran, J., Brenner, T. J. and DiNardo, S. (2000). Somatic control over the germline stem cell lineage during Drosophila spermatogenesis. Nature 407, 754–757.

Vara, C., Paytuvi-Gallart, A., Cuartero, Y., Le Dily, F., Garcia, F., Salva-Castro, J., Gomez, H. L., Julia, E., Moutinho, C., Aiese Cigliano, R., et al. (2019). Three-Dimensional Genomic Structure and Cohesin Occupancy Correlate with Transcriptional Activity during Spermatogenesis. Cell Rep 28, 352–367 e359.

Venken, K. J., Popodi, E., Holtzman, S. L., Schulze, K. L., Park, S., Carlson, J. W., Hoskins, R. A., Bellen, H. J. and Kaufman, T. C. (2010). A molecularly defined duplication set for the X chromosome of Drosophila melanogaster. Genetics 186, 1111–1125.

Vermeulen, K., Van Bockstaele, D. R. and Berneman, Z. N. (2003). The cell cycle: a review of regulation, deregulation and therapeutic targets in cancer. Cell Prolif 36, 131–149.

Voog, J., Sandall, S. L., Hime, G. R., Resende, L. P., Loza-Coll, M., Aslanian, A., Yates, J. R., 3rd, Hunter, T., Fuller, M. T. and Jones, D. L. (2014). Escargot restricts niche cell to stem cell conversion in the Drosophila testis. Cell Rep 7, 722–734.

Wang, Z. and Lin, H. (2005). The division of Drosophila germline stem cells and their precursors requires a specific cyclin. Curr Biol 15, 328–333.

Wang, Z. A. and Kalderon, D. (2009). Cyclin E-dependent protein kinase activity regulates niche retention of Drosophila ovarian follicle stem cells. Proc Natl Acad Sci U S A 106, 21701–21706.

Wharton, W. (1983). Hormonal regulation of discrete portions of the cell cycle: commitment to DNA synthesis is commitment to cellular division. J Cell Physiol 117, 423–429.

Yamaguchi, M., Hayashi, Y. and Matsukage, A. (1995). Essential role of E2F recognition sites in regulation of the proliferating cell nuclear antigen gene promoter during Drosophila development. J Biol Chem 270, 25159–25165.

Zappia, M. P. and Frolov, M. V. (2016). E2F function in muscle growth is necessary and sufficient for viability in Drosophila. Nat Commun 7, 10509.

Zappia, M. P., Rogers, A., Islam, A. and Frolov, M. V. (2019). Rbf Activates the Myogenic Transcriptional Program to Promote Skeletal Muscle Differentiation. Cell Rep 26, 702–719 e706.

