## Supplementary figures for "Cell cycle exit and stem cell differentiation are coupled through regulation of mitochondrial activity in the Drosophila testis"

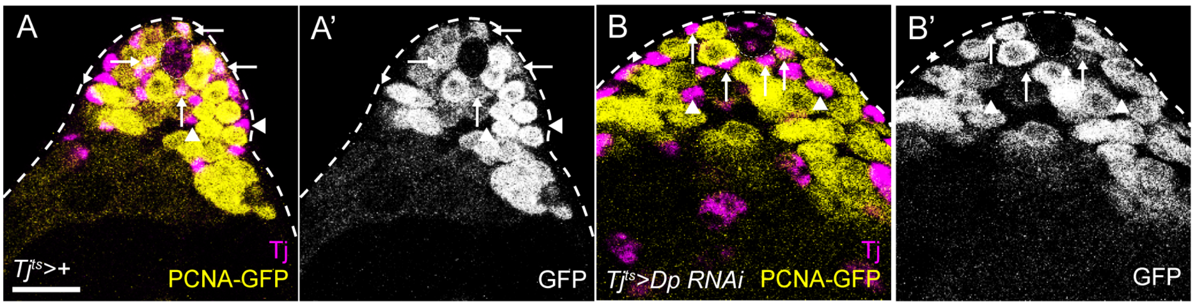


**Figure S1. E2f1/Dp activity is detected in the somatic lineage.** Expression of the canonical E2f1/Dp reporter *PCNA-GFP* (yellow, single channels in A’,B’) in the testis. Tj (magenta) labels CySCs and early cyst cells. In controls (A), GFP expression is detected in Tj-positive CySCs adjacent to the hub (arrows), while no expression is present in differentiated cyst cells (arrowhead). (B) Dp knockdown in the somatic lineage results in decreased expression in Tj-positive cells around the hub (arrows), indicating that PCNA-GFP expression reports on endogenous E2f1/Dp activity. Dotted lines outline the hub. Scale bars: 20µm.

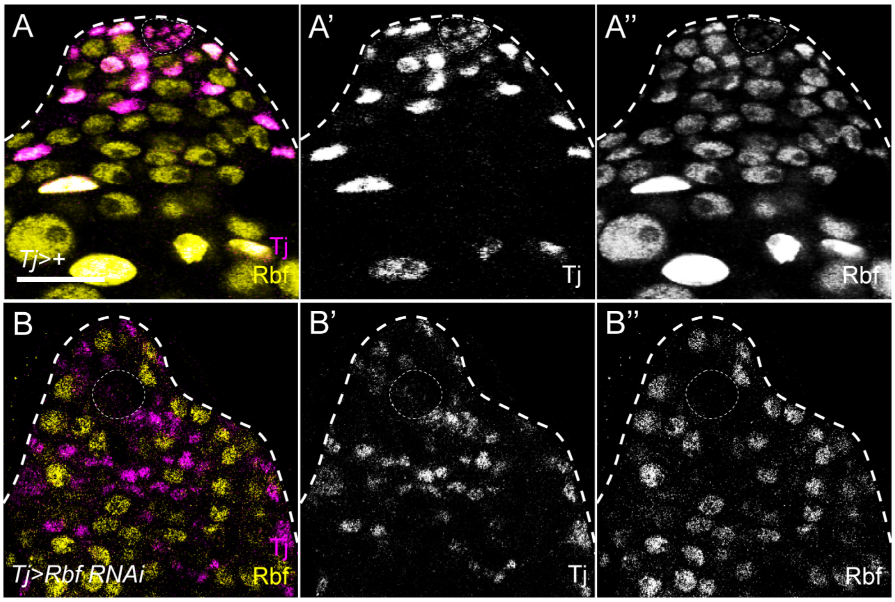


**Figure S2. Rbf is expressed in CySCs and efficiently knocked down by RNAi.** Testes labelled with antibodies against Rbf (yellow, single channels in A”,B”) and Tj (magenta, single channels in A’,B’). In controls (A), Rbf is detected in all cells in the Drosophila testis, including CySCs adjacent to the hub (dotted line) and differentiating cyst cells. Expression increases in cyst cells distant from the hub. Rbf is also present in germ cells and weakly in hub cells. (B) Expression of Rbf RNAi with *tj-Gal4* results in a complete lack of any detectable Rbf protein in Tj-positive cells. Dotted lines outline the hub. Scale bars: 20µm.

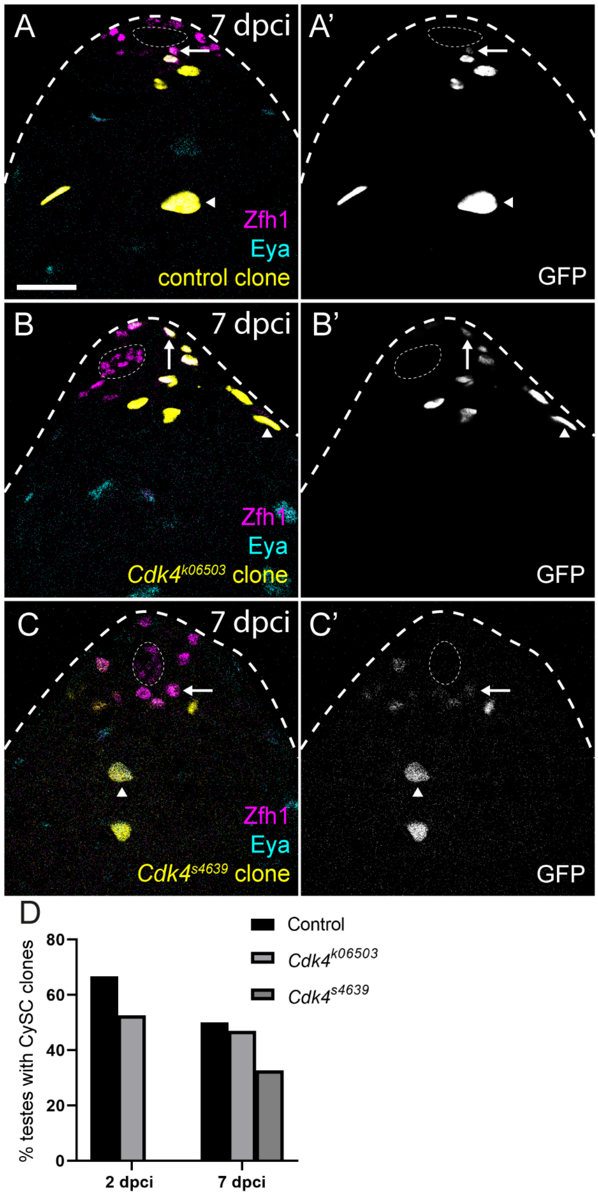


**Figure S3. *Cdk4* is not required for CySC self-renewal.** Positively-marked clones labelled by GFP expression (yellow, single channels in A',B',C') were generated and recovered at 7 days post clone induction (dpci). Zfh1 (magenta) identifies CySCs and early daughters, and Eya (cyan) marks differentiated cyst cells. Control clones (A) at 7 dpci contained both Zfh1-expressing CySCs (arrow) and Eya-positive cyst cells (arrowhead). Clones mutant for independent alleles of *cdk4* (B,C) were similarly recovered at 7 dpci and contained Zfh1-expressing CySCs (arrows), indicating that *Cdk4* is not necessary for CySC self-renewal. (D) Quantification of clone recovery rates, showing the fraction of testes containing controls and *Cdk4* mutant CySC clones. Mutant clone recovery rates were not significant different from controls, determined by Fisher's exact test. See Table 1 for n values. Dotted lines outline the hub. Scale bars: 20µm.

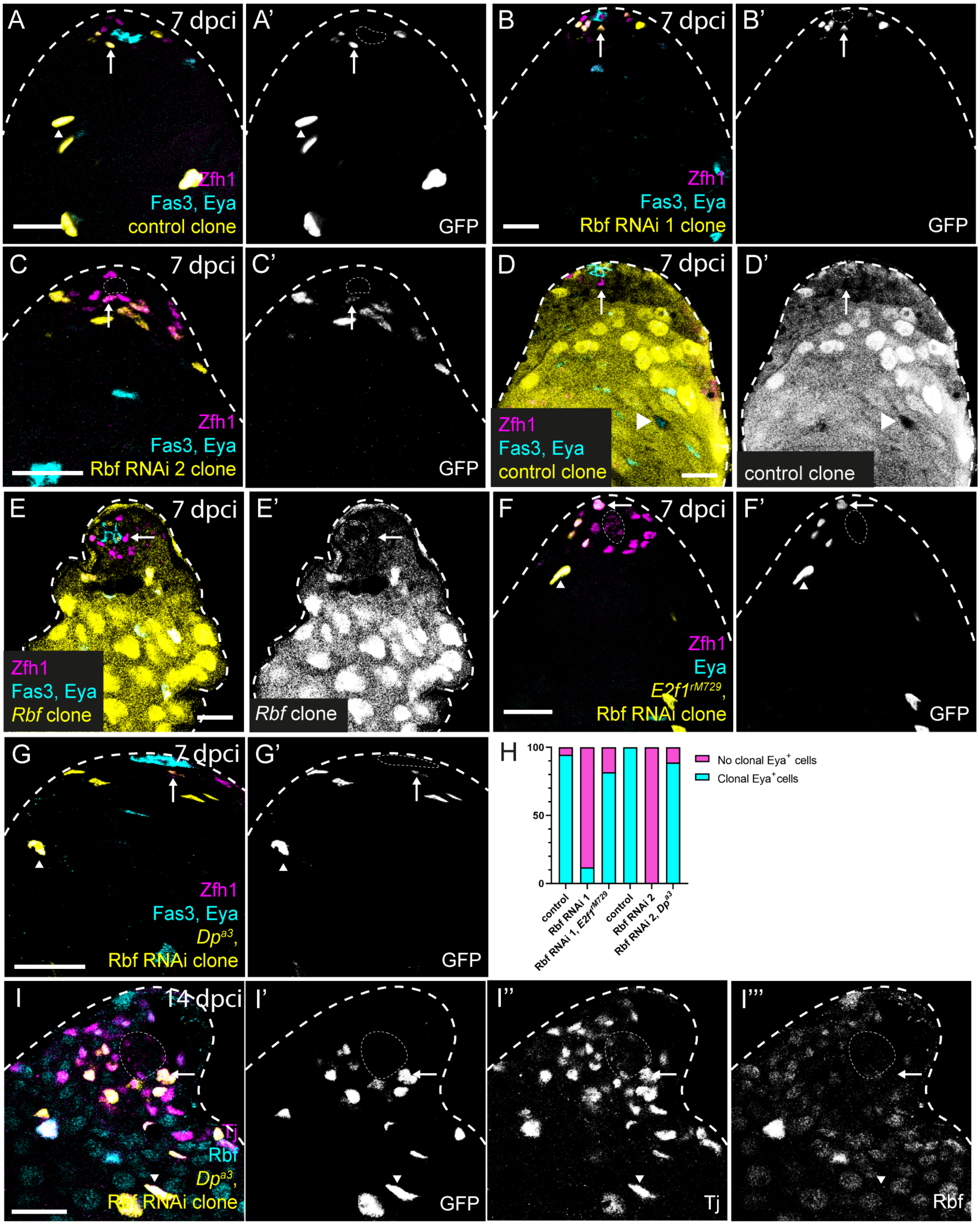


**Figure S4. Rbf inhibition of E2f1/Dp is required autonomously for cyst cell differentiation.** (A-C) CySC clones expressing GFP (yellow, single channels in A',B',C'). Zfh1 (magenta) marks CySCs and early daughters, Eya marks differentiated cyst cells and Fas3 marks the hub. Control clones at 7 dpci (A) are recovered adjacent to the hub (arrow) and contain differentiated cyst cells (arrowhead). (B,C) Clones expressing two separate RNAi constructs targeting Rbf consisted exclusively of Zfh1-expressing cells and contained no Eya-expressing cyst cells. (D,E) Negatively-marked clones lacking a duplication rescuing *Rbf*, identified by lack of GFP expression (yellow, single channel in D',E'). See Methods for information about genotypes. Control clones are composed of Zfh1-positive CySCs (arrow) and Eya-positive cyst cells (arrowhead) at 7 dpci. (E) In contrast, *Rbf* hemizygous mutant clones contained only Zfh1-expressing cells (arrow), indicating that the *Rbf* mutant phenotype in CySCs is similar to RNAi knockdown. (F-I) Differentiation of Rbf RNAi clones is rescued by loss-of-function of *E2f1* or *Dp*. Clones mutant for *E2f1* (F) or *Dp* (G) in which Rbf RNAi was expressed were composed of both Zfh1-expressing CySCs (arrows) and Eya-positive differentiated cyst cells (arrowheads). (H) Quantification of clones containing Eya-positive cells in the different genotypes, showing the rescue of *Rbf* loss-of-function by *E2f1* or *Dp*. (I) Rbf protein (yellow, single channel in I'") is absent from *Dp* mutant clones expressing Rbf RNAi. Cyst lineage clones are identified by GFP expression (yellow, single channel I') and Tj expression (magenta, single channel I"). Dotted lines outline the hub. Scale bars: 20µm.

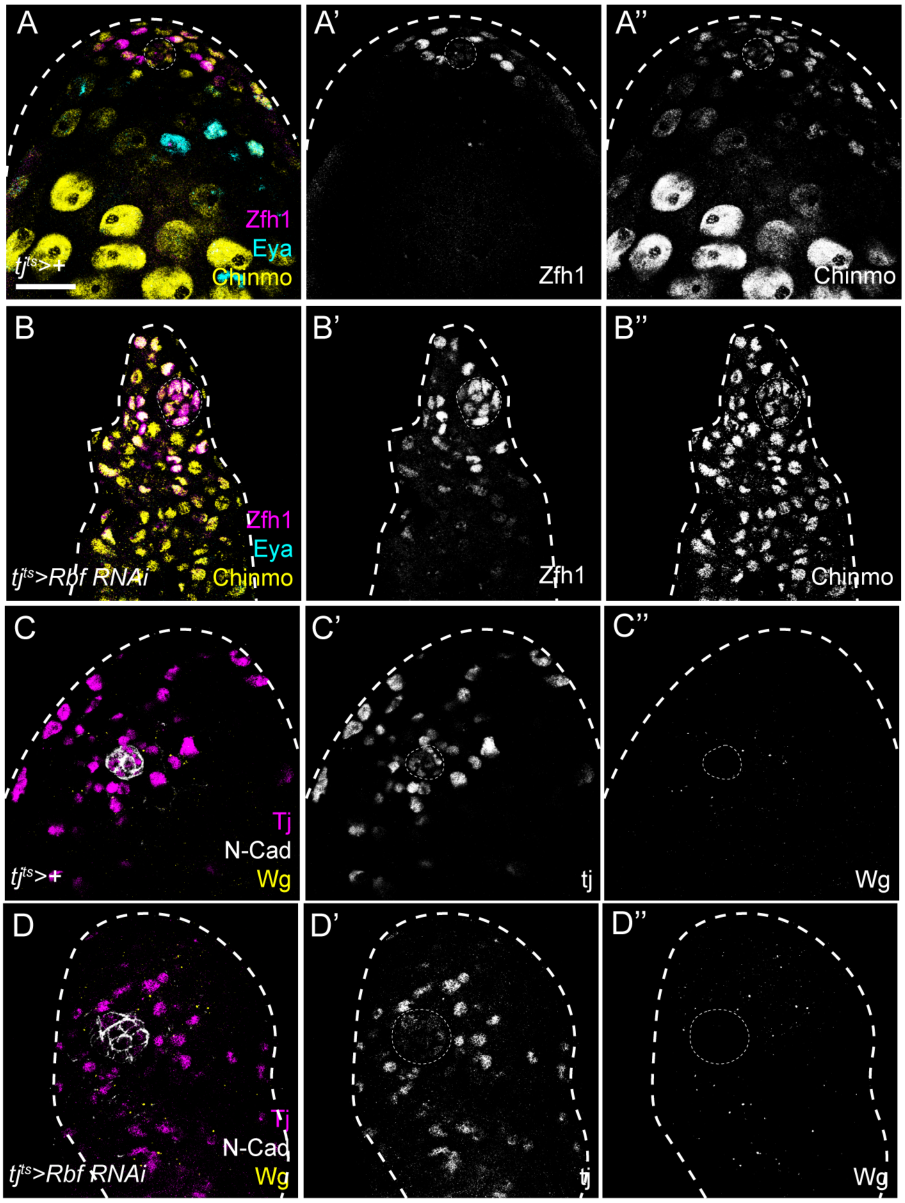


**Figure S5. Rbf-deficient cells express markers of CySCs.** (A,B) Testes labelled with antibodies against Chinmo (yellow, single channel in A",B"), Zfh1 (magenta, single channel in A',B') to mark CySCs and early daughters, and Eya (cyan), to mark differentiated cyst cells. In controls (A), Chinmo is expressed in the hub, Zfh1-positive CySCs, GSCs, identified as cells adjacent to the hub negative for Zfh1 expression, and late germ cells. Expression is absent from Eya-positive cyst cells. When Rbf was knocked down in somatic cells with *tj*$^{ts}$, Chinmo expression was expanded and detected in Zfh1-positive somatic cells distant from the hub. Expression of Chinmo in unlabelled early germ cells was also expanded. (C,D) Wg expression (yellow, single channels in C",D") in testes labelled with antibodies against N-Cadherin (N-Cad, grey) to label the hub and Tj (magenta, single channels in C',D') to label CySCs and early cyst cells. In controls, Wg puncta are visible around somatic cells close to the hub. Wg staining was detected far from the hub in testes with somatic Rbf knockdown, indicating an expansion of undifferentiated CySC-like cells. Dotted lines outline the hub. Scale bars: 20µm.

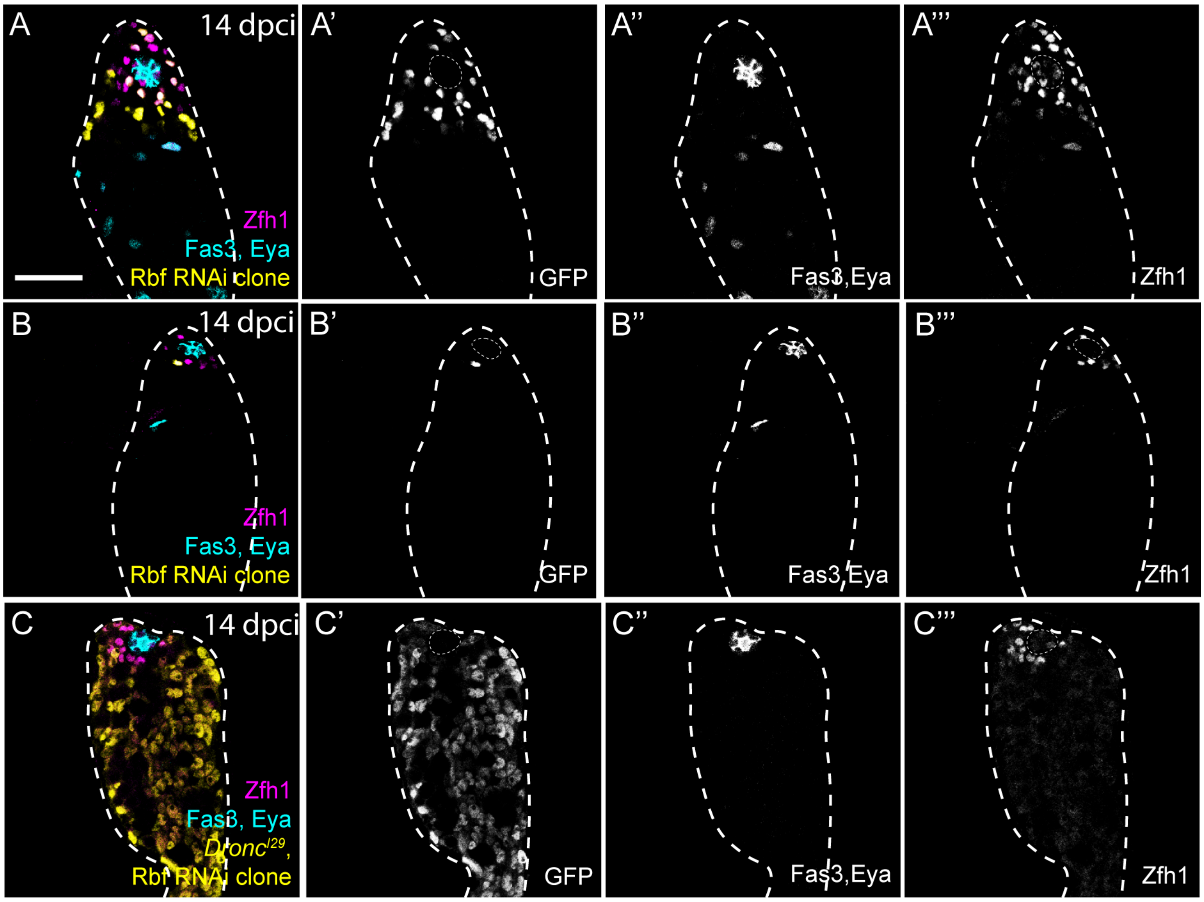


**Figure S6. Apoptosis limits the proliferation of Rbf-deficient CySCs, but is not responsible for the lack of differentiation.** CySC clones at 14 dpci positively-labelled with GFP expression (yellow, single channel in A',B',C'). Fas3 (cyan, single channel in A'',B'',C'') labels the hub, Zfh1 (magenta, single channel in A''',B''',C''') marks CySCs and early daughter cells, and Eya (cyan, single channel in A'',B'',C'') marks differentiated cyst cells. Expression of Rbf RNAi in otherwise wild type clones (A,B) resulted in clones composed only of Zfh1-positive cells and lacking Eya-positive differentiated cyst cells. While most clones contained many cells (A), indicative of proliferation, occasional clones were observed that were composed of very few cells (B). (C) Rbf RNAi expression in clones also mutant for *Dronc* resulted in large clones but these were still devoid of Eya-expressing cyst cells, indicating that cell death restricts the over-proliferation of Rbf-deficient cells but is not responsible for their inability to differentiate. Dotted lines outline the hub. Scale bars: 20µm.

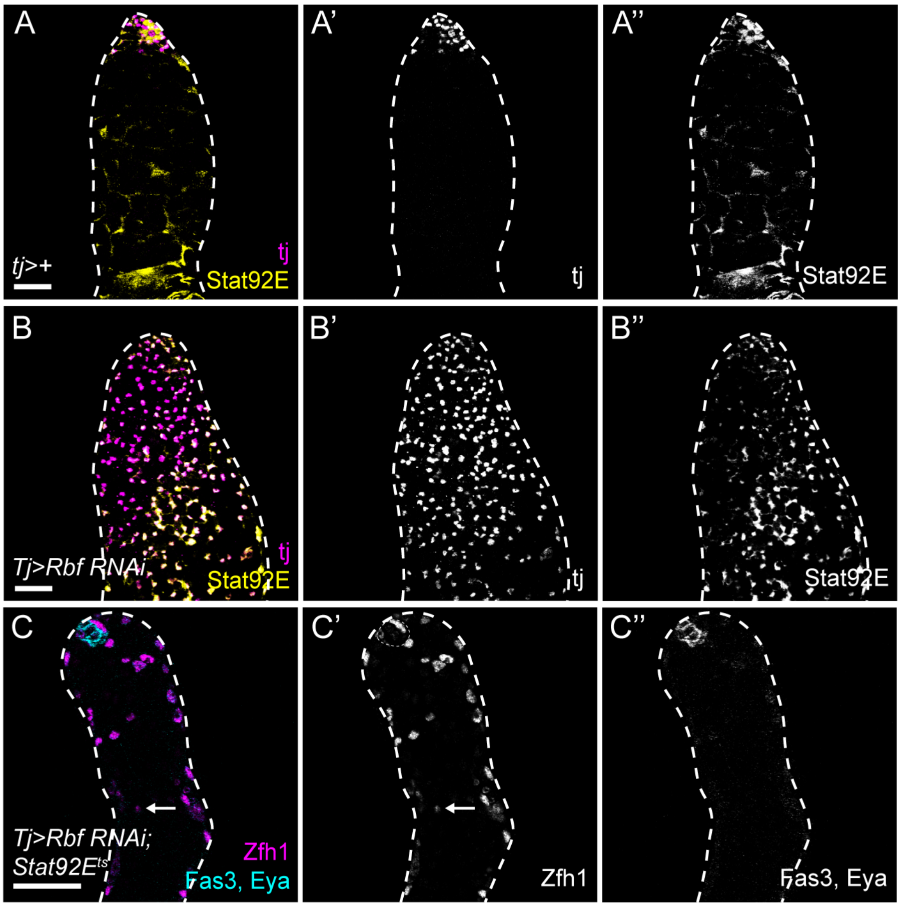


**Figure S7. Increased JAK/STAT signalling does not mediate the expansion of CySC-like cells caused by Rbf knockdown.** (A,B) Testes labelled with antibodies against Stat92E (yellow, single channels in A",B"), which is stabilised where JAK/STAT signalling is active, and Tj (magenta, single channels in A',B') to mark CySCs and early cyst cells. In controls (A), Stat92E is detected around the hub. In testes somatically depleted for Rbf (B), stabilised Stat92E is present in Tj-positive cells throughout the testes. (C) Rbf knockdown in the somatic lineage in a temperature-sensitive *Stat92E* mutant (*Stat92E*$^{ts}$) raised for 10 days at the restrictive temperature results in expansion of Zfh1 (magenta, single channel in C') away from the hub (arrow) and a lack of Eya-positive cells (cyan, single channel in C"), similar to Rbf knockdown in control animals. Dotted lines outline the hub. Scale bars: XXµm in A,B; 20µm in C.

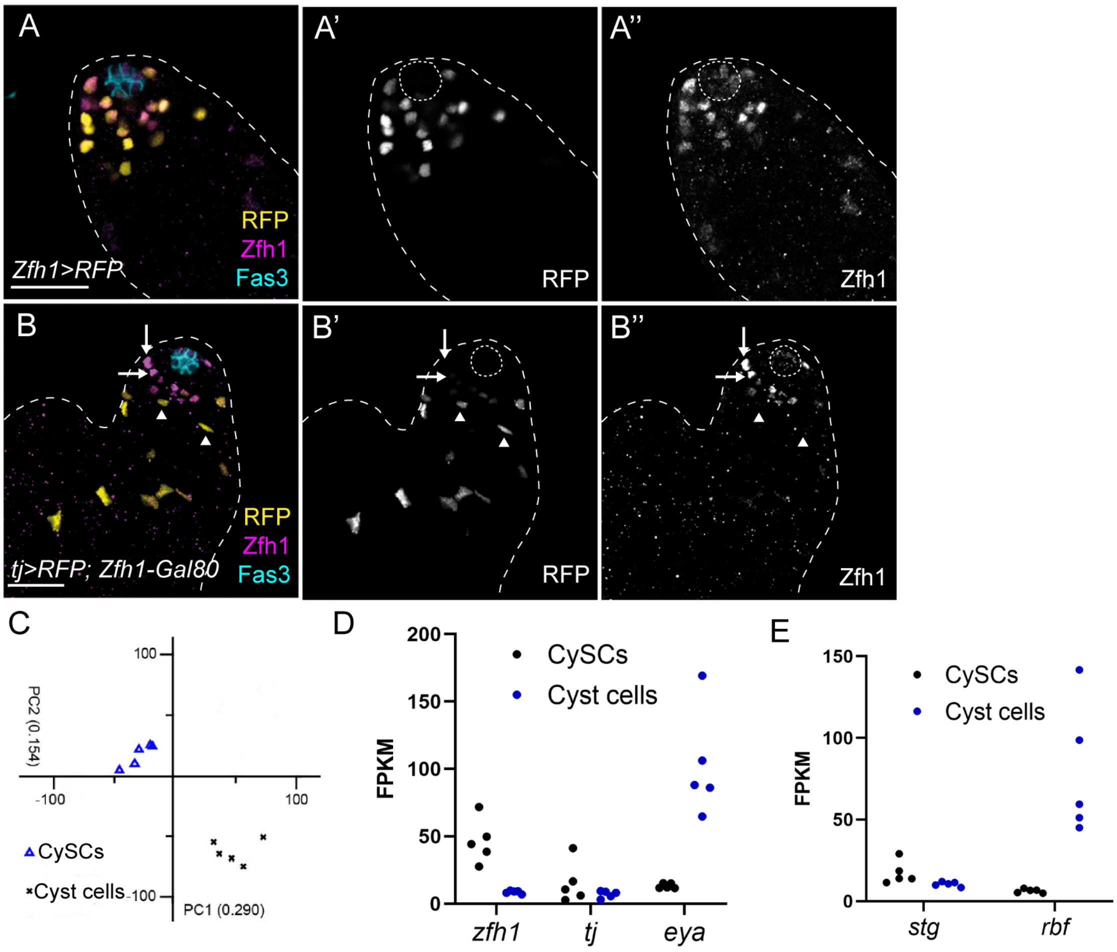


**Figure S8. Effective isolation and transcriptome determination of CySCs and cyst cells.** (A) Expression of RFP (yellow, single channel A’) in *Zfh1>RFP* testes to isolate CySCs. Presence of RFP correlates with Zfh1 expression (magenta, single channel A”). The hub is labelled with Fas3 (cyan). (B) Expression of RFP (yellow, single channel in B’) in *tj>RFP; Zfh1-Gal80* testes to isolate differentiating cyst cells. Zfh1-positive cells (magenta, single channel in B”) close to the hub (Fas3, cyan) do not express RFP (arrows), whereas early differentiating cells (arrowheads) negative for Zfh1 express RFP. (C) Principal component analysis of the transcriptomes of five biological replicates of sorted cell populations showing that CySCs and cyst cells cluster separately. (D) Individual values for reads (in Fragments Per Kilobase of transcript per Million mapped reads (FPKM)) for each replicate of genes encoding known markers of CySCs and cyst cells in the sorted CySC population (black) and cyst cells (blue). As expected, CySCs express higher levels of *zfh1* and *tj*, whereas early cyst cells express higher levels of *eya*. (E) Individual values for reads in FPKM of genes encoding cell cycle components. *Stg* expression which was enriched in CySCs (black) relative to cyst cells (blue), while *Rbf* was upregulated in cyst cells.

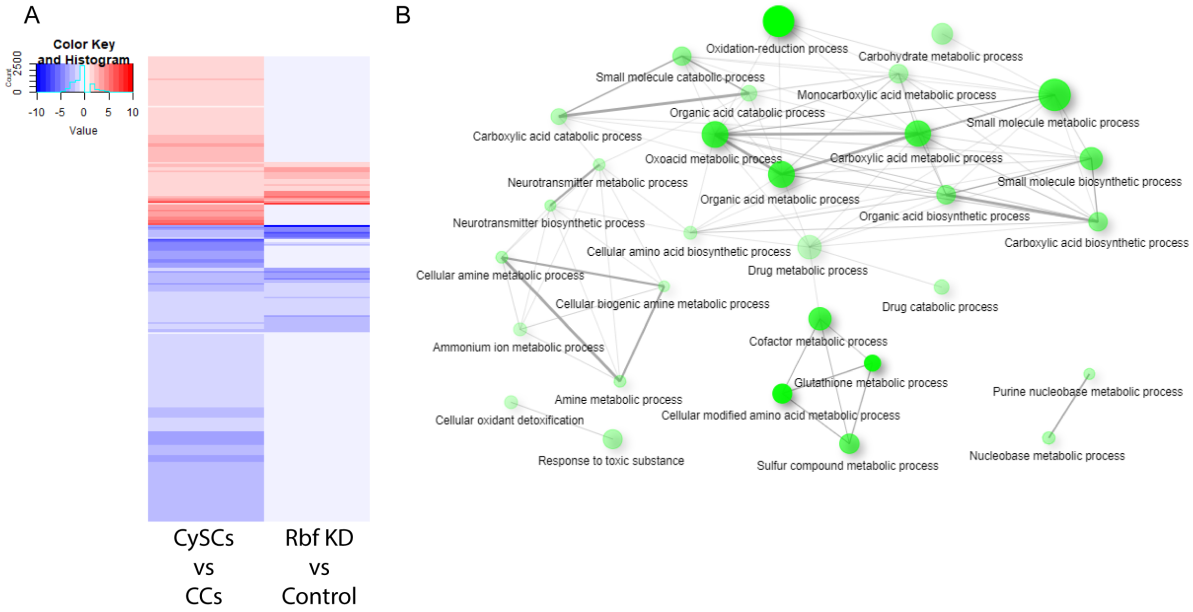


**Figure S9. Rbf-deficient cells show similar gene expression profiles to CySCs.** (A) Heatmap representing the shared gene differential expression among experiments. Differentially-expressed genes between CySCs and cyst cells (left column) were used as a reference. Genes more highly expressed in CySCs relative to cyst cells are shown in red and genes enriched in cyst cells are shown in blue. Changes for these genes in Rbf knockdown compared to controls are shown in the right column and colour-coded according to differential expression in Rbf knockdowns. Pale blue in the right column indicates genes differentially expressed between control CySCs and cyst cells that were not differentially expressed when Rbf-knockdown somatic cells were compared to controls. Approximately 27% of genes are differentially expressed in both experiments, and show similar valence of expression change. (B) Gene Ontology analysis of genes downregulated in both control CySCs compared to cyst cells and Rbf-deficient cells compared to control. The size of the circles represents the number of genes in each biological process. A higher significance of enrichment is shown as stronger colouring of the circle. Lines connect nodes with at least 15% of shared genes and their thickness increases with the percentage of shared genes. Shared biological processes include carbohydrate metabolism and oxidative metabolism and oxidation-reduction balance, suggesting that somatic knockdown of Rbf results in a similar metabolic gene expression profile to CySCs.

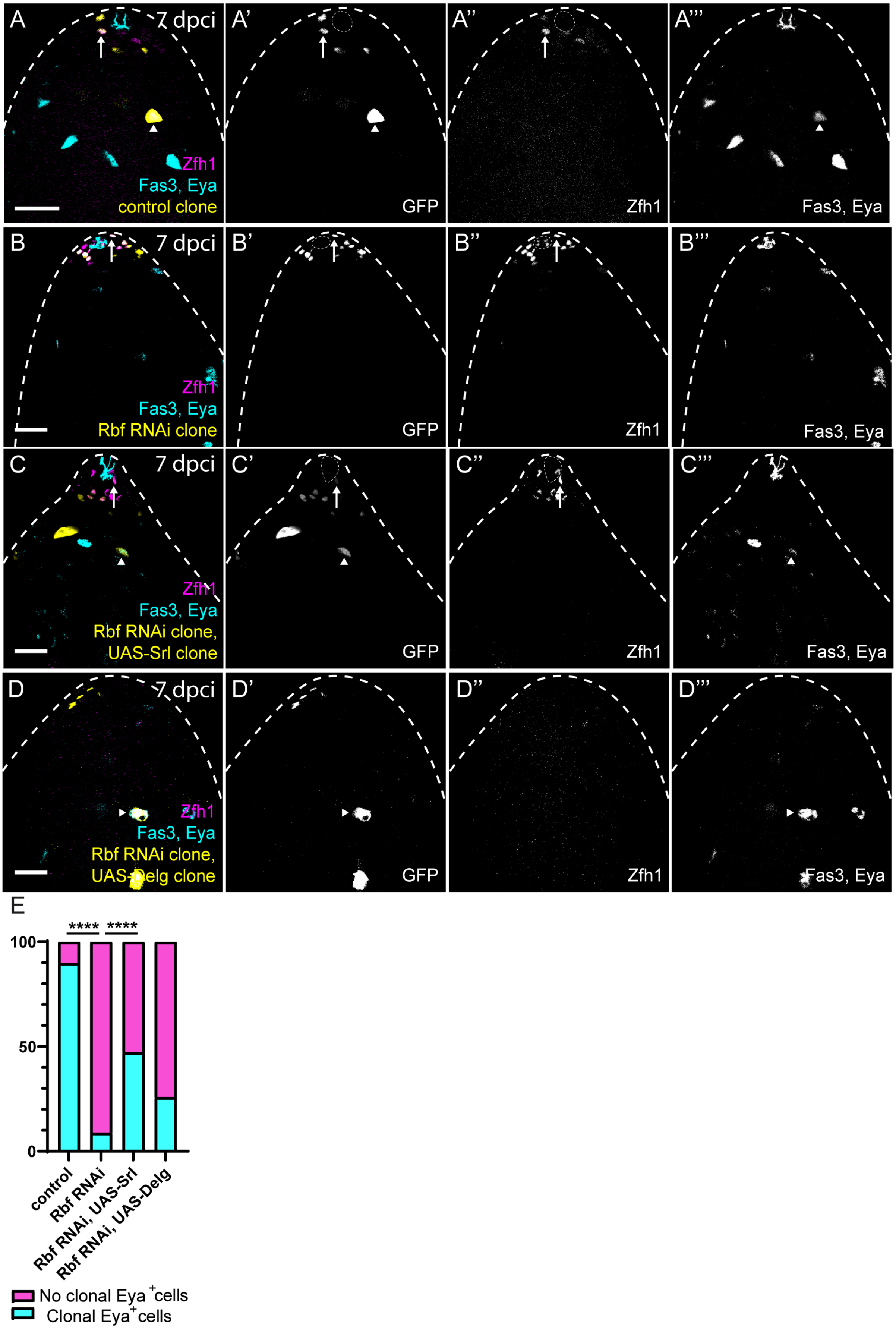

A
7 dpci
Zfh1
Fas3, Eya
control clone
A'
GFP
A''
Zfh1
A'''
Fas3, Eya
B
7 dpci
Zfh1
Fas3, Eya
Rbf RNAi clone
B'
GFP
B''
Zfh1
B'''
Fas3, Eya
C
7 dpci
Zfh1
Fas3, Eya
Rbf RNAi clone,
UAS-Srl clone
C'
GFP
C''
Zfh1
C'''
Fas3, Eya
D
7 dpci
Zfh1
Fas3, Eya
Rbf RNAi clone,
UAS-Delg clone
D'
GFP
D''
Zfh1
D'''
Fas3, Eya
E
****
****
100
50
0
control
Rbf RNAi
Rbf RNAi, UAS-Srl
Rbf RNAi, UAS-Delg
No clonal Eya+ cells
Clonal Eya+ cells

**Figure S10. Differentiation is rescued by co-expression of Srl or Delg upon clonal loss of Rbf.** (A-D) Positively-labelled clones marked by GFP expression (yellow, single channels in A'-D'). Zfh1 (magenta, single channel in A"-D") marks CySCs and early daughters, while Eya (cyan, single channel in A'"-D'") marks differentiated cyst cells. The hub is labelled with Eya (cyan). Control clones (A) contained both Zfh1-positive CySCs (arrow) and Eya-positive cyst cells (arrowhead), while only Zfh1-expressing cells were detected in Rbf RNAi-expressing clones (B, arrow). Co-expression of Srl (C) or Delg (D) with Rbf RNAi resulted in clones that contained Eya-positive cells (arrowheads). In D, the hub is in an adjacent plane. (E) Graph showing the percentage of clones containing Eya-positive cells (cyan bars). Only 9% of Rbf RNAi-expressing clones contained Eya-positive cells (n=49 for control, n=46 for Rbf RNAi, $P<0.0001$ compared to control clones, Fisher's exact test). Co-expression of Srl resulted in a significant rescue compared to Rbf RNAi alone to 47% (n=36, $P<0.0001$), while 26% of clones co-expressing Delg contained Eya-positive cells, approaching statistical significance (n=35, P=0.06). Dotted lines outline the hub. Scale bars: 20µm.
