## Supplementary text for "Cell cycle exit and stem cell differentiation are coupled through regulation of mitochondrial activity in the Drosophila testis"

**Apoptosis does not account for the lack of differentiation in Rbf knockdown.**

We ruled out the possibility that Rbf-deficient differentiated cyst cells were lost to apoptosis by knocking down *Rbf* in CySC clones mutant for *Dronc*, the initiator caspase-9 homologue (Domingos and Steller, 2007). At 14 dpci, *Dronc* mutant clones lacking Rbf did not contain any Eya-expressing cells (Fig. S6). In otherwise wild type Rbf-knockdowns, we observed occasional clones composed of fewer than 10 cells (4/33 clones); these small clones were hardly ever observed in *Dronc* mutant clones in which Rbf was knocked down (Fig. S6C) (1/25 clones), indicating that apoptosis may limit the proliferation of Rbf-deficient CySCs, but cannot account for the lack of differentiated cyst cells.

**JAK/STAT signalling is not required for ectopic CySCs induced by Rbf knockdown.**

We asked whether increased self-renewal signalling could be responsible for the expansion of stem-like cells downstream of Rbf knockdown. JAK/STAT is the main self-renewal pathway in CySCs and gain-of-function in the cyst lineage results in over-proliferation of CySCs and a block in differentiation similar to *Rbf* loss-of-function (Issigonis et al., 2009; Kiger et al., 2001; Leatherman and Dinardo, 2008). Indeed, we detected increased stabilisation of the sole Drosophila STAT, Stat92E, upon Rbf knockdown (Fig. S7A,B). However, knocking down Rbf in the cyst lineage in a temperature-sensitive *Stat92E* mutant resulted in expansion of the Zfh1-expressing population, and lack of Eya-expressing differentiated cyst cells, similar to Rbf knockdown alone (Fig. S7C). Thus, Rbf loss-of-function results in ectopic CySC-like cells independently of JAK/STAT signalling.

**Supplementary references.**

**Domingos, P. M. and Steller, H.** (2007). Pathways regulating apoptosis during patterning and development. *Curr Opin Genet Dev* **17**, 294-299.

**Issigonis, M., Tulina, N., de Cuevas, M., Brawley, C., Sandler, L. and Matunis, E.** (2009). JAK-STAT signal inhibition regulates competition in the Drosophila testis stem cell niche. *Science* **326**, 153-156.

**Kiger, A. A., Jones, D. L., Schulz, C., Rogers, M. B. and Fuller, M. T.** (2001). Stem cell self-renewal specified by JAK-STAT activation in response to a support cell cue. *Science* **294**, 2542-2545.

**Leatherman, J. L. and Dinardo, S.** (2008). Zfh-1 controls somatic stem cell self-renewal in the Drosophila testis and nonautonomously influences germline stem cell self-renewal. *Cell Stem Cell* **3**, 44-54.
